# A multifunctional Wnt regulator underlies the evolution of coat pattern in African striped mice

**DOI:** 10.1101/2022.12.12.520043

**Authors:** Matthew R. Johnson, Sha Li, Christian F. Guerrero-Juarez, Pearson Miller, Benjamin J. Brack, Sarah A. Mereby, Charles Feigin, Jenna Gaska, Qing Nie, Jaime A. Rivera-Perez, Alexander Ploss, Stanislav Y. Shvartsman, Ricardo Mallarino

## Abstract

Animal pigment patterns are excellent models to elucidate mechanisms of biological organization. Although theoretical simulations, such as Turing reaction-diffusion systems, recapitulate many animal patterns, they are insufficient to account for those showing a high degree of spatial organization and reproducibility. Here, we compare the coats of the African striped mouse (*Rhabdomys pumilio*) and the laboratory mouse (*Mus musculus*) to study the molecular mechanisms controlling stripe pattern formation. By combining transcriptomics, mathematical modeling, and mouse transgenics, we show that *Sfrp2* regulates the distribution of hair follicles and establishes an embryonic prepattern that foreshadows pigment stripes. Moreover, by developing and employing *in vivo* gene editing experiments in striped mice, we find that *Sfrp2* knockout is sufficient to alter the stripe pattern. Strikingly, mutants also exhibit changes in coat color, revealing an additional function of *Sfrp2* in regulating hair color. Thus, a single factor controls coat pattern formation by acting both as an orienting signaling mechanism and a modulator of pigmentation. By uncovering a multifunctional regulator of stripe formation, our work provides insights into the mechanisms by which spatial patterns are established in developing embryos and the molecular basis of phenotypic novelty.

---

The skin of many vertebrate species displays characteristic periodic pigment patterns, such as spots and stripes, which play key roles in mediating intra-and inter-specific communication^1–3^. Because of their visual accessibility, extreme diversity, and widespread occurrence among multiple species, color patterns represent a fascinating model to understand the mechanisms underlying biological organization. While this research has been mostly restricted to a handful of traditional laboratory model species, recent advances in genomics and experimental approaches now offer the possibility of studying wild-derived species to uncover the molecular, cellular, and developmental processes that have generated the astonishing diversity of color patterns found in nature.

Theoretical simulations involving Turing reaction-diffusion systems, in which activators and inhibitors interact to establish organized spatial patterns, can closely recapitulate a wide variety of periodic color patterns seen in nature^4–6^. Notably, however, Turing-generated patterns are variable and sensitive to stochastic perturbation.^7^. That is, Turing mechanisms are insufficient, by themselves, to explain the formation of periodic color patterns seen in many species, which are characterized by a high degree of spatial organization and reproducibility across individuals^8–12^. Therefore, a long-standing challenge has been to uncover the molecular, cellular, and developmental events by which periodic color patterns are specified and organized.

The African striped mouse (*Rhabdomys pumilio*), a rodent that exhibits a naturally evolved coat pattern of dark and light parallel stripes (**Fig. 1a**), is a useful model in which to explore molecular mechanisms underlying periodic color patterns because this mouse can be maintained and reared in the lab, allowing for controlled experiments and development of molecular tools^13^. Moreover, the African striped mouse and the laboratory mouse (*Mus musculus*), the premier model species in mammalian/skin research, are closely related^13^, opening the door for powerful comparative studies. Here, we use a variety of multidisciplinary approaches to uncover the developmental mechanisms controlling coat pattern formation in striped mice.

**Fig. 1.**
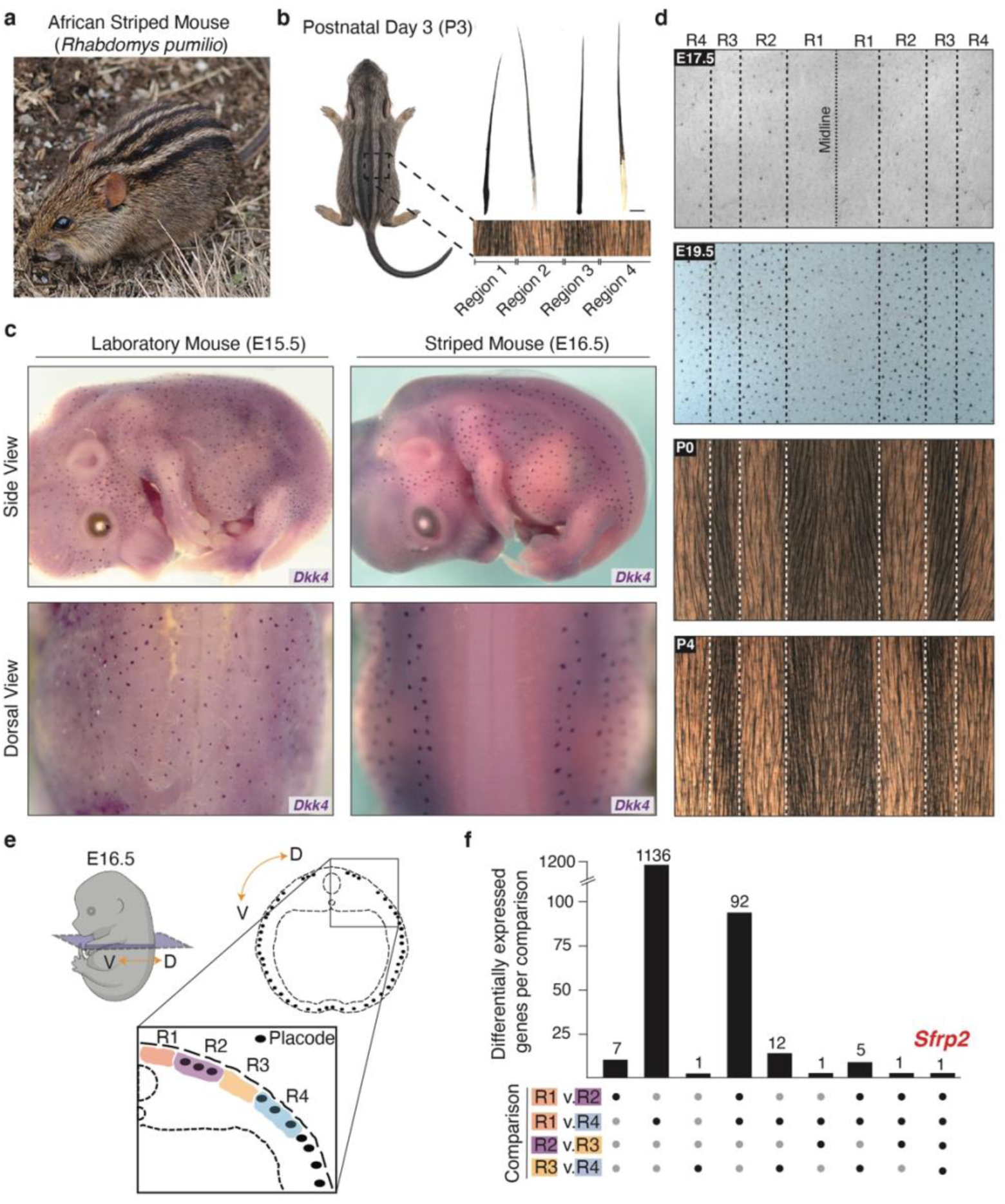
Development of hair placodes and pigment patterns in the African striped mouse. **a**, An adult striped mouse in the field. **b**, Postnatal Day 3 (P3) striped mouse pup displaying the characteristic striped pattern and corresponding differences in hair length across dorsal regions. **c**, Side and dorsal views of Embryonic Day 15.5 (E15.5) laboratory mouse and E16.5 striped mouse embryos displaying spatial patterns of hair placode formation, as visualized by whole-mount in situ hybridization for Dkk4. **d**, Light-microscopy images of striped mouse flat mount skins from developmental and postnatal stages (E16.5 to P4) showing how patterns of hair placode development foreshadow pigmentation stripes. **e**, Schematic of an E16.5 striped mouse embryo displaying regions (R1-R4) dissected for bulk-level RNA-seq analysis. **f**, UpSet plot summarizing differentially expressed genes (DEGs) (padj < 0.05) between R1-R4. Each row of dots represents a single pairwise comparison. Black dots indicate there were DEGs between a given comparison (number of DEGs plotted as columns) whereas grey dots indicate there were not. Photo credit in **a**: Trevor Hardaker

## Striped mice display regional variation in embryonic placode formation

Striped mice exhibit differences in hair length between striped skin regions that are apparent at early postnatal stages (**Fig. 1b**)^10^. We hypothesized that these differences may reflect stripe-specific alterations in the developmental timing of hair follicle placodes (hereafter referred to as placodes). In laboratory mice (*Mus musculus*), placodes are visible at embryonic day 15.5 (E15.5) and are evenly distributed throughout the dorsal skin (**Fig. 1c**)^14, 15^. Remarkably, this stereotyped spatial pattern differs considerably in striped mice. At the equivalent embryonic stage (i.e., the emergence of visible placodes, ∼E16.5 in striped mice), whole mount *in situ* hybridizations for early markers of placode formation (e.g., *Dkk4, Ctnnb1, Wnt10b, Wif1, Eda,* and *Dkk1*)^16^ revealed that striped mouse dorsal placodes develop in a stripe-like, spatially restricted manner, whereby they are present in some regions along the dorsal skin but absent in others (**Fig. 1c and Extended Data Fig. 1a, b**). Approximately two days later, at E18.5, placodes eventually become visible in regions previously devoid of them, as evidenced by thickening of the epidermis and the appearance of molecular markers (**Extended Data Fig. 2a, b**). Notably, stripe-like dorsal areas where placodes fail to form initially correspond to regions that will constitute the eventual dark pigment stripes, explaining why hair from dark stripes is shorter at birth and suggesting that spatially restricted patterns of embryonic placode formation foreshadow pigmentation stripes (**Fig. 1d**)^10^.

**Fig. 2.**
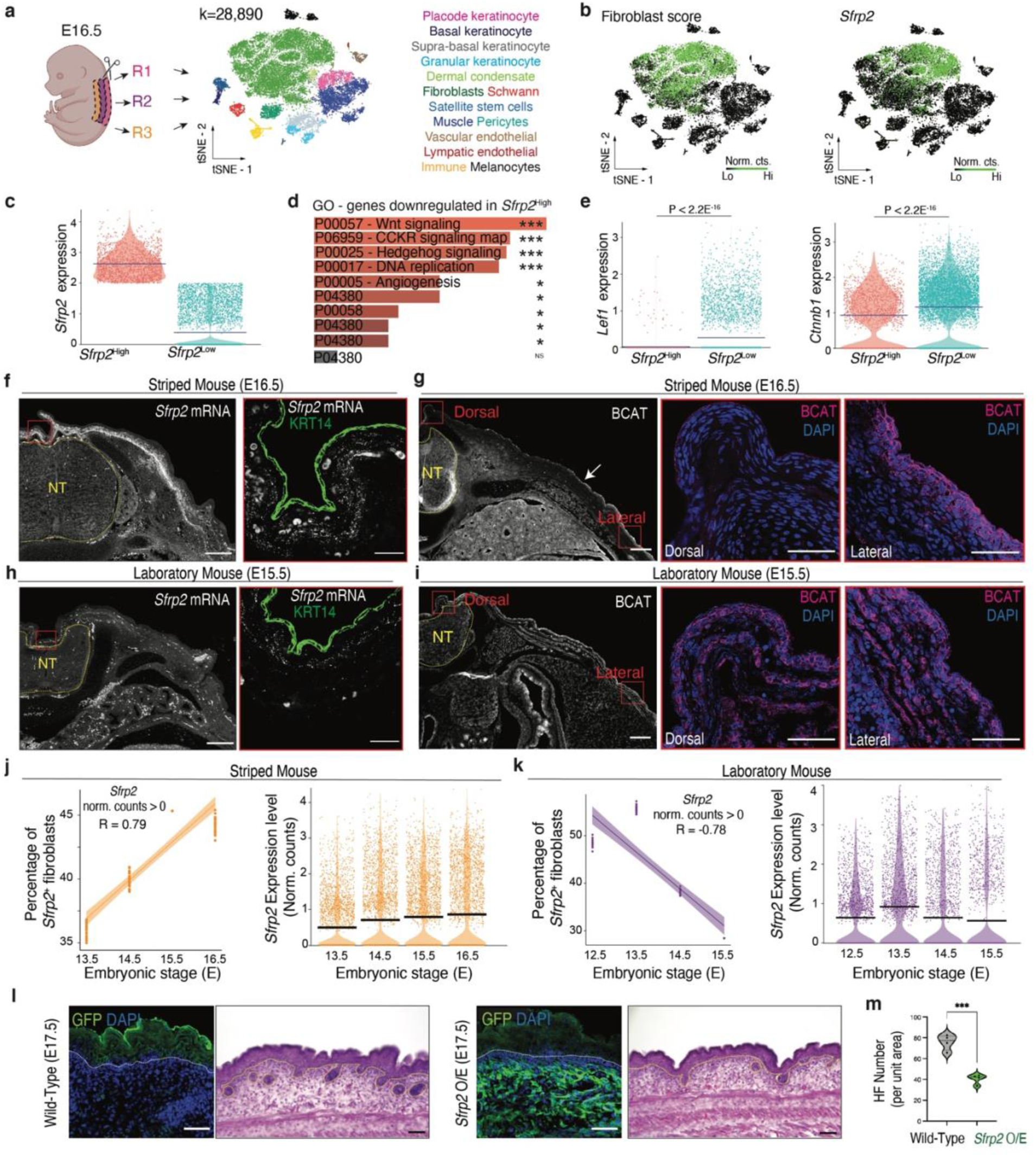
Relationship between *Sfrp2* and Wnt signaling in embryonic skin. **a**, Schematic of skin regions (R1-R3) dissected for scRNA-seq (left) and cell-type clustering (right). **b**, Dermal fibroblasts (green, left) cluster with *Sfrp2* expressing cells (green, right). **c**, Fibroblasts sorted by high (> 2 UMI) and low (< 2 UMI) *Sfrp2* expression. **d**, Ontology analysis of genes downregulated in *Sfrp2*^High^ cells, show Wnt signaling as the main enriched pathway (Enrichr, Padj < 0.05). **e**, *Lef1* and *Ctnnb1* show upregulation in *Sfrp2*^low^ cells (non-parametric Wilcoxon Rank Sum test). **f-g**, *In situ* hybridization for *Sfrp2* coupled with immunofluorescence (IF) for KRT14 in stage-matched striped mouse and laboratory mouse embryos. **h-i**, IF for CTNNB1 in stage-matched striped mouse and laboratory mouse embryos. Red boxes denote zoomed-in regions. **j-k**, Comparative scRNA-seq analysis shows both the percentage of fibroblasts that express *Sfrp2* (left, Spearman’s Correlation, R = 0.79) and the expression levels of *Sfrp2* within striped mouse fibroblasts (**j**) increase in the days prior to visible placode formation while, in laboratory mice (**k**), both the percentage of fibroblasts expressing *Sfrp2* (left, Spearman’s correlation, R = -0.78) and *Sfrp2* expression levels within those fibroblasts (right) decrease in the days prior to visible placode formation. **l-m**, Representative GFP and H&E stained transverse sections showing the distribution of SFRP2 and hair follicles in control and transgenic mice (**l**) as well as differences in hair follicle number (**m**) (N = 4; *** P < 0.001). Scale bars: 200µm (zoomed out) and 25µm (zoomed in) in (**f** and **h**); 200µm (zoomed out) and 50µm (zoomed in) in (**g** and **i**); and 50µm in (**j**). NT = neural tube.

### Sfrp2 is expressed in a dorsoventral gradient during placode formation

To identify regulators of the striped mouse placode developmental pattern, we performed an unbiased bulk-level RNA-sequencing (RNA-seq) screen for genes differentially expressed between placode rich and placode barren skin regions in E16.5 embryos. We used the boundaries between these regions as a proxy to mark and isolate four distinct dorsal skin regions: R1 (the dorsalmost, placode barren region), R2 (the dorsalmost, placode rich region), R3 (the ventralmost, placode barren region), and R4 (the ventralmost, placode rich region) (**Fig. 1e**). Next, we performed a series of pairwise comparisons (R1 vs. R2; R1 vs. R4; R2 vs. R3; and R3 vs. R4) to identify differentially expressed genes between placode barren (R1, R3) and placode rich (R2, R4) regions. Although each pairwise comparison yielded several differentially expressed genes (**Fig. 1f and Supplementary Table 1**), only one gene, the Wnt modulator *Sfrp2*, was differentially expressed in all four comparisons. Surprisingly, however, *Sfrp2* was neither up nor downregulated in placode barren (R1, R3) or placode rich regions (R2, R4). Rather, this gene was always upregulated in the dorsalmost region among the two being compared (i.e., R1>R2, R1>R4, R2>R3, and R3>R4).

### Sfrp2 negatively regulates Wnt signaling in striped mouse embryonic skin

Since Wnt signals are required for the initiation of hair follicle development (*16*), *Sfrp2* constituted an attractive candidate for patterning striped mouse placodes. However, as *Sfrp2* can have dual roles as both an activator and repressor of Wnt signaling, depending on cellular microenvironment^17–20^, the relationship between *Sfrp2* and Wnt signaling in striped mouse skin remained unclear. Moreover, whether *Sfrp2* plays a role in placode formation has not been previously investigated. To study this, we dissociated dorsal skin from regions R1, R2, and R3 of E16.5 embryos, as before, and performed single-cell 3’ -RNA-sequencing (scRNA-seq, 10X Genomics). We combined an equal number of cells from R1-R3 to generate a single pool representative of striped mouse E16.5 dorsal skin and identified fourteen distinct cell types based on expression of known molecular markers (**Fig. 2a and Extended Data Table 1**)^21–23^. *Sfrp2* was specifically expressed in a subset of dermal fibroblasts, as judged by co-localization with canonical fibroblast markers (**Fig. 2b and Extended Data Table 1**). Next, we clustered cells based on *Sfrp2* expression to define a population of *Sfrp2*-high (i.e., > 2UMIs) and *Sfrp2*-low (i.e., < 2 UMIs) fibroblasts (**Fig. 2c**). Differential gene expression analysis between *Sfrp2*-high and *Sfrp2*-low cells yielded numerous genes (Padj <0.05), 482 of which were upregulated and 272 that were downregulated in *Sfrp2*-high compared to *Sfrp2*-low cells (**Supplementary Table 2**). Among the genes upregulated in *Sfrp2*-low fibroblasts, Gene Ontology analysis identified regulation of Wnt signaling as the most significantly enriched pathway (**Fig. 2d**). Indeed, expression of numerous Wnt-related genes, including key activators of the pathway (i.e., *Lef1* and *Ctnnb1*), were higher in *Sfrp2-*low (**Fig. 2e and Supplementary Table 2**) compared to *Sfrp2*-high cells, demonstrating that *Sfrp2* and Wnt activation are negatively correlated in striped mice dermal fibroblasts.

To confirm and extend our previous observation, we examined spatial patterns of *Sfrp2* expression and Wnt signaling markers. RNA *in situ* hybridizations in cross sections from E16.5 striped mice embryos, coupled to immunohistochemistry (IHC) for an epidermal marker (KRT14), indicated that expression of *Sfrp2* was strong in mesenchymal cells directly above the neural tube and decreased laterally, a result consistent with our bulk-level RNA-seq analysis (**Fig. 2f**). At this same stage, immunofluorescence (IF) for LEF1 and CTNNB1 showed that these markers were readily detectable in epidermal cells from lateral and ventral regions of the embryo but were absent from cells above the neural tube and adjacent areas, where *Sfrp2* expression was highest (**Fig. 2g and Extended Data Fig. 3**). This pattern contrasted with what was seen in stage-matched laboratory mouse embryos (E15.5), which had overall lower levels of dermal *Sfrp2* expression (**Fig. 2h**) and uniform levels of LEF1 and CTNNB1 throughout the epidermis, including above the neural tube (**Fig. 2i and Extended Data Fig. 3**).

**Fig. 3.**
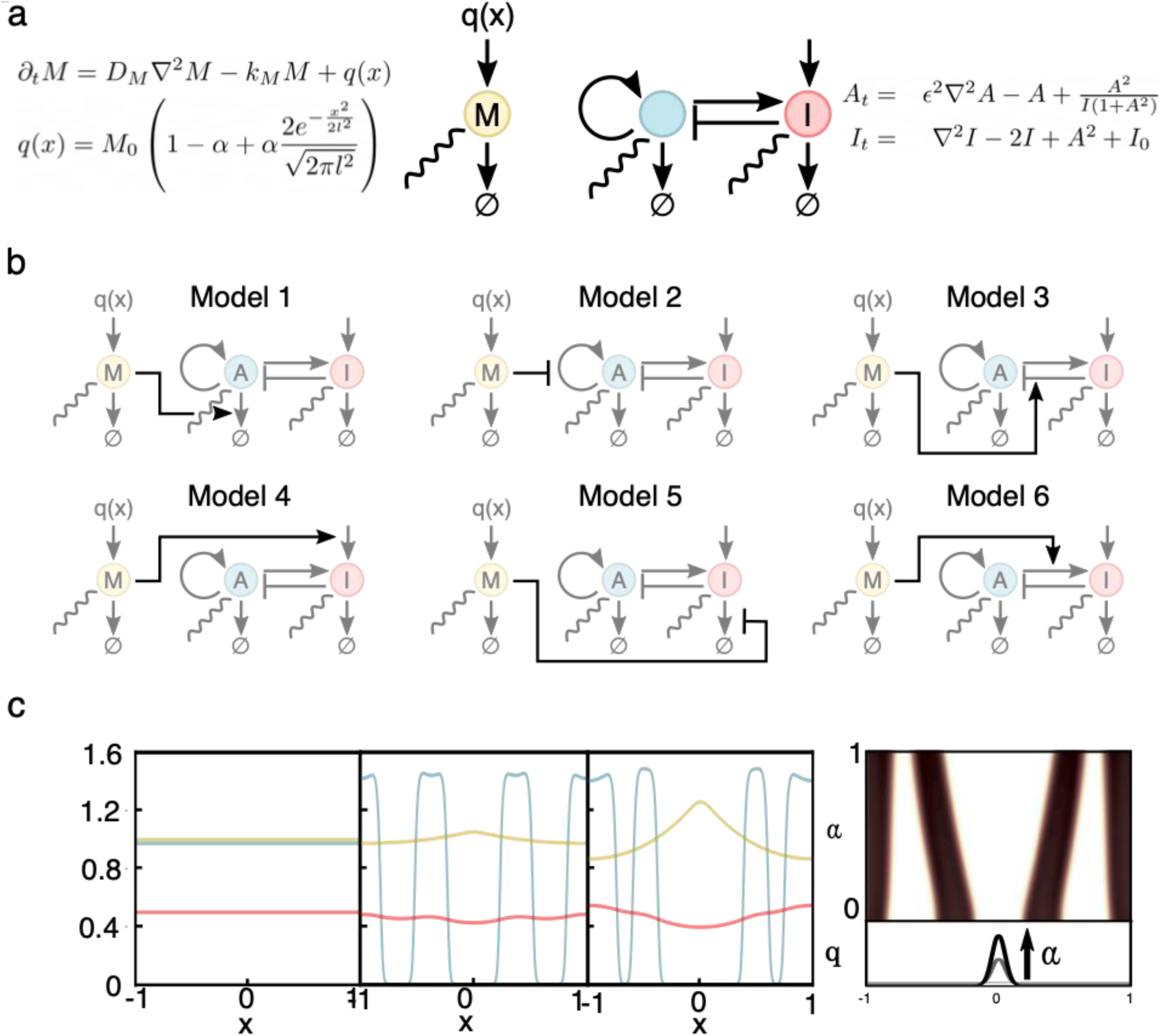
Modulator gradients control stripe patterning in a reaction-diffusion system. **a**, We present a two module explanator model for stripe formation. The left module describes a freely diffusible modulator *M* produced by a non-uniform source term *q(x).* The non-uniformity of modulator expression is controlled by the gradient steepness parameter *α*. The right module is a variant of the canonical Gierer-Meinhardt model for Turing pattern formation featuring saturated activator production, which is known to reliably produce stripe patterns. **b**, We consider a range of possible couplings between these two modules, all of which produce qualitatively equivalent results. For further details, see **Extended Data** Fig. 4. **c**, Example results from model variant 1: (Left) In the absence of a modulator gradient, the stripe-less/uniform state is stable. Introduction of modulator gradients destabilize the uniform state and establish stripes (Middle, Right). **d**, Decreasing *α* (i.e., decreasing the strength of the modulator gradient) corresponds to a narrowing of the medial white region and widening of the lateral white regions (i.e., regions corresponding placode-less regions in E16.5 striped mice embryos). This effect is independent of the model choice.

Notably, the marked differences in *Sfrp2* expression between striped mouse and laboratory mouse were not only evident during placode formation, but also in stages prior. Specifically, as indicated by scRNAseq, the number of *Sfrp2* expressing fibroblasts, as well as the average *Sfrp2* expression levels within those fibroblasts, steadily increased in the days preceding striped mouse placode emergence (E13.5-E15.5) (**Fig. 2j**). This pattern starkly contrasted with laboratory mouse, where *Sfrp*2 expressing fibroblasts dramatically decreased over the same period (E12.5-E14.5) and average *Sfrp2* expression levels within fibroblasts did not change (**Fig. 2k**).

Taken together, our scRNA-seq data coupled to RNA *in situ*/IF analysis indicate that *Sfrp2* and Wnt signaling are negatively correlated, suggest that dermal *Sfrp2* expression acts to inhibit Wnt signaling, and indicate that the observed patterns are unique to striped mouse.

### Sfrp2 overexpression alters placode formation in laboratory mouse

Wnt signaling is a primary determinant of hair follicle formation and spacing, as perturbation of members of this pathway leads to alterations in hair follicle number and distribution^15, 16, 24, 25^. This raises the intriguing hypothesis that *Sfrp2* may be acting to directly regulate striped mouse placode development. If this is the case, one would predict that increasing expression of *Sfrp2* in embryonic skin with low endogenous levels of *Sfrp2*, such as that of laboratory mouse (**Fig. 2f, h, j, k**)^22, 23^, should alter placode formation. To test this, we generated *Rosa^Sfrp^*^2^*^-GFP^* mice, a strain that allowed us to activate *Sfrp2* transcription in the presence of a tissue-specific Cre driver. To establish whether dermal expression of *Sfrp2* was sufficient to regulate placode formation, as implied by our striped mouse data, we crossed *Rosa^Sfrp^*^2^*^-GFP^* mice with *Dermo-Cre* mice, a mesenchymal driver of Cre^26, 27^. Indeed, double-transgenic embryos (*Dermo-Cre*;*Rosa^Sfrp^*^2^*^-GFP^*) analyzed at E17.5 showed a significant reduction in the number of placodes/hair follicles, compared to control littermates (N= 4) (**Fig. 2l, m**). Thus, our transgenic experiments in laboratory mice demonstrate a functional relationship between *Sfrp2* and embryonic placode development in which elevated levels of *Sfrp2* in the dermis leads to a marked reduction in the number of placodes/hair follicles.

### Modeling implicates Sfrp2 in patterning striped mouse placodes

During striped mouse mid-embryogenesis, *Sfrp2* is expressed in a dorsoventral gradient and acts to negatively regulate Wnt signaling. Moreover, in line with its role as a Wnt inhibitor, *Sfrp2* caused a reduction in placode formation. The dorsoventral expression pattern of a placode inhibitor was, *a priori*, unexpected. However, previous work has pointed to upstream morphogen gradients as a means for establishing stripe-like patterns^28^. In lab mouse, placode spacing is regulated by a self-organizing Turing reaction-diffusion system in which activators and inhibitors interact to establish an organized spatial pattern^29^. While the classic picture of stripe formation via a Turing instability features the emergence of patterns in response to global, spatially uniform perturbations, recent theoretical work has increasingly emphasized the capacity of spatial gradients to trigger the emergence of stripes^30^. We therefore turned to mathematical modelling to understand how a dorsoventral gradient could influence an underlying Turing pattern. Our approach was to start with a canonical set of equations for biochemical pattern formation, the Gierer-Meinhardt (GM) model^31^, and separately a simple model of a diffusible modulator with spatially non-uniform expression (**Fig. 3a**). As a modulator can be coupled to the GM system in various ways, we constructed a family of model variants which we studied in parallel (e.g., in model 1, the modulator accelerates the local degradation of the activator while in model 2, it interferes with the strength of the activator’s positive feedback loop, etc.) (**Fig. 3b**). Our analysis focused on two related phenomena: (1) the destabilization of the uniform steady state by the introduction of modulation, and (2) the relationship between modulation and stripe spacing.

The bifurcation structure of GM is well established in the absence of modulation, allowing us to choose a starting point in our parameter space featuring a stable uniform steady state and stable stripe patterns^32, 33^. For our initial assay, we altered the parameter controlling non-uniformity of modulator production. In doing so, we found that for each of our model variants, a sufficiently strong gradient in modulator production would destabilize the uniform steady state, leading to the appearance of stripes (**Fig. 3c and Extended Data Fig. 4**). Thus, by altering the underlying Turing activity (i.e., changing it from uniform to non-uniform) a dorsoventral modulator may indeed explain differences between the absence (e.g., laboratory mouse) or presence (e.g., striped mouse) of stripes.

**Fig. 4.**
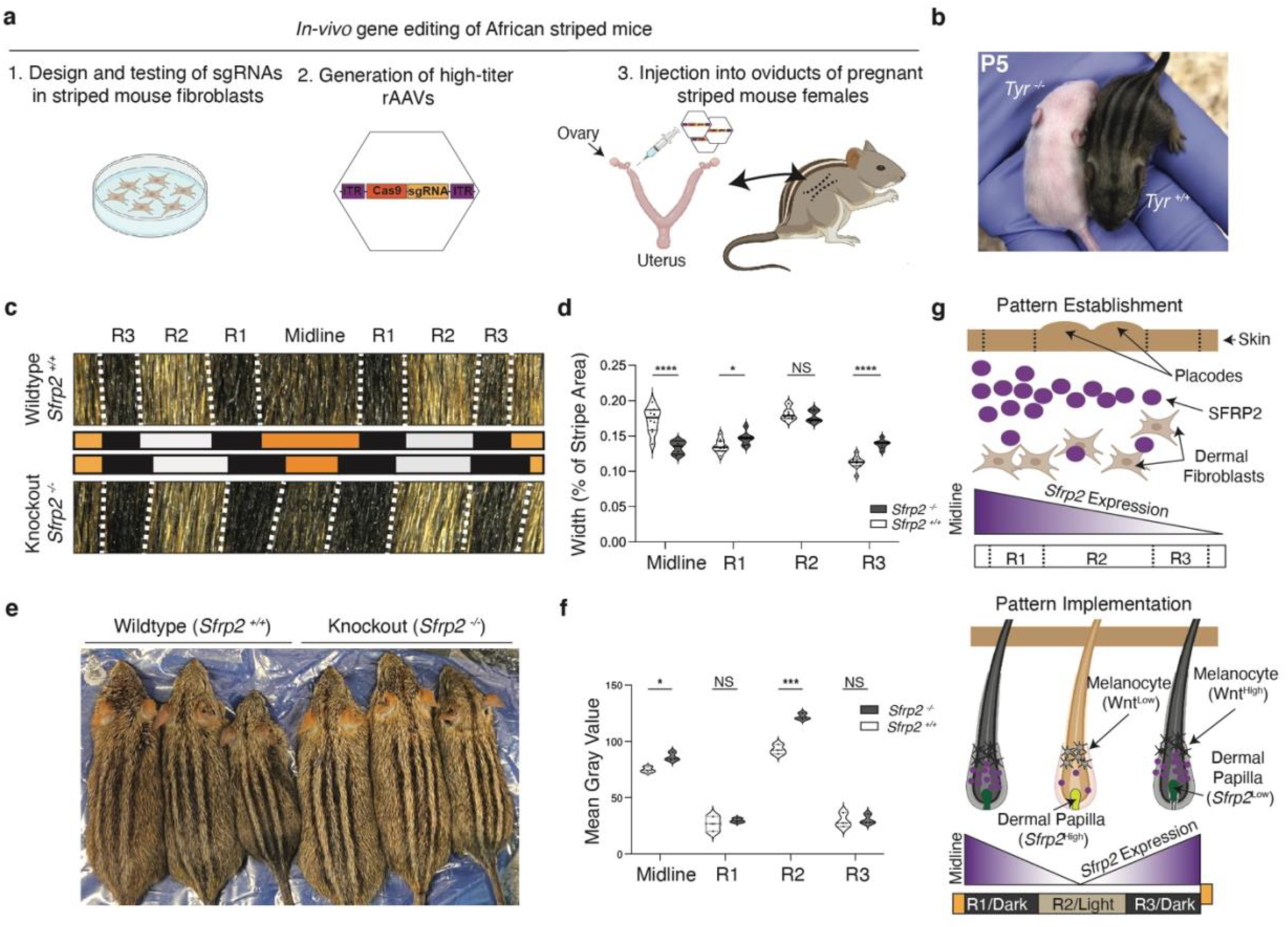
*In vivo* genome editing reveals that *Sfrp2* regulates striped mouse coat patterns. **a**, Schematic of the in vivo gene editing method used. **b**, Postnatal Day 5 (P5) Tyrosinase wildtype (Tyr^+/+^) and knockout (Tyr^-/-^) striped mouse littermates. **c-d**, *Sfrp2* regulates stripe pattern width. Shown are representative images (**c**) and measurements (**d**) revealing differences in stripe width between wildtype (*Sfrp2*^+/+^) and *Sfrp2* knockout (*Sfrp2*^-/-^) pups. Statistical significance in (**d**) (**** P < 0.0001; *** P < 0.001; ** P < 0.01; *P < 0.05; *Sfrp2*^+/+^, N = 10; *Sfrp2*^-/-^, N = 7) was assessed using an ANOVA test. **e**, Representative differences in coat color between *Sfrp2*^+/+^ and *Sfrp2*^-/-^ adult mice taken as a single image. **f**, Quantification of pigmentation between *Sfrp2*^+/+^ and *Sfrp2*^-/-^ adult mice reveal stripe-specific changes in color. Statistical significance in panel (**f**) (*** P < 0.001; *P < 0.05; *Sfrp2*^+/+^, N = 3; *Sfrp*2^-/-^, N = 3). **g**, Model describing the role of *Sfrp2* in establishing (top) and implementing (bottom) the coat pattern in striped mice.

We next used additional simulations to study the relationship between alterations to the modulator gradient and potential consequences on stripe spacing. Our results demonstrated the continued stability of stripes, regardless of the magnitude of the modulator gradient (**Extended Data Fig. 4b**). More strikingly, however, we found that altering the strength of the modulator gradient led to stereotyped changes in the spacing of the stripes: in all models, changing only the magnitude of the modulator gradient produced subtle changes in stripe width (**Fig. 3c, d and Extended Data Fig. 4b**).

Taken together, our mathematical simulations coupled to our experimental data implicate *Sfrp2* as a key component of dorsoventral gradient that organizes placode patterns in striped mouse. Furthermore, our simulations make testable predictions about the phenotypic changes that result from altering such a gradient.

### In vivo gene editing in striped mice leads to changes in coat pattern

To establish a direct causal link between *Sfrp2* and placode formation patterns as well as test the predictions from our mathematical simulations, it is critical to perform functional experiments directly in striped mice. To this end, we adapted a CRISPR-based experimental strategy in which recombinant adeno-associated viruses (rAAVs) are used to deliver CRISPR-CAS9 reagent into the oviduct of pregnant females. This *in vivo* gene editing technique is based on the ability of rAAVs to transduce the zona pellucida on intact pre-implantation embryos^34^ (**Fig. 4a**). First, to establish the feasibility of using this approach for inducing gene knockouts in striped mice, we injected pregnant females carrying pre-implantation embryos with an rAAV packaged with Nme2Cas9 and a single-guide RNA (sgRNA) expression cassettes directed to the coding sequence of *Tyrosinase* (*Tyr*), the rate limiting enzyme for melanin production^35, 36^. Visual inspection of pups coupled to genotyping revealed this approach resulted in efficient gene editing, enabling us to generate complete knockout animals (**Fig. 4b**). To our knowledge, this is the first time that *in vivo* gene editing has been performed in a wild-derived mammalian species.

Encouraged by our proof-of-principle results, we next designed multiple sgRNA’s against the 5’ coding region of *Sfrp2* (**Extended Data Fig. 5A**). We then tested each sgRNA in striped mouse immortalized dermal fibroblasts and chose the one that had the highest indel-inducing efficiency. Next, we injected an rAAV containing Nme2Cas9 and sg*Sfrp2* expression cassettes into the oviducts of plugged females. From a total of 21 females injected, we obtained 57 pups, 14 of which carried gene edits in *Sfrp2*. After mating F0 founders and wildtype animals (*Sfrp2^+/+^*) to ensure germline transmission, we crossed F1 heterozygous animals (*Sfrp2^+/-^*) to generate F2 *Sfrp2* knockout (*Sfrp2^-/-^*) mice. We used western blot to confirm the successful elimination of SFRP2 protein (**Extended Data Fig. 5b**). Analysis of coat phenotypes from early postnatal (P3) stages revealed significant and consistent differences in the coat patterns of F2 *Sfrp2^-/-^*and *Sfrp2^+/+^* striped mice: we observed significant changes in the relative width of the stripes between *Sfrp2^+/+^* and *Sfrp2^-/-^* mice, as predicted by our models (**Fig. 4c, d** and **Fig. 3b**). Thus, by knocking out *Sfrp2*, the dorsoventral organizing gradient is perturbed, effectively altering the boundaries of the striped mouse’s coat pattern.

### Sfrp2 regulates pigmentation via activation of Wnt signaling

In addition to changes in stripe pattern, we noticed that mutant animals had obvious differences in coat coloration, with *Sfrp2^-/-^*mice showing overall lighter pigmentation than *Sfrp2^+/+^* mice (**Fig. 4e, f**). Previous studies have shown that pigment production/melanogenesis can be controlled by Wnt signals secreted from the dermal papilla^37, 38^, a specialized mesenchymal component of the hair follicle that signals to pigment-producing cells, or melanocytes. Thus, to explore a potential link between *Sfrp2* and melanogenesis, we analyzed the skin of striped mice at P4, a period in which pigment is being actively synthesized and deposited in the growing hair shaft. Fluorescent *in situ* hybridizations on skin sections revealed that *Sfrp2* was specifically expressed in the dermal papilla (**Extended Data Fig. 6a**). Next, to test the effect of *Sfrp2* expression on melanocytes, we stably transduced an immortalized mouse melanocyte cell line (Melan-A) with a lentiviral vector carrying the *Sfrp2* cDNA and used qPCR to measure the expression of *Tyr* and *Mitf*, two genes that promote melanin synthesis^35, 39^. Compared to melanocytes transduced with a control virus, cells overexpressing *Sfrp2* showed a significant increase in *Tyr* and *Mitf* (**Extended Data Fig. 6b**). Moreover, they also had marked increases in Wnt target genes, including *Axin2*, *C-myc*, and *Ccnd1*^40–42^. Notably, these results are consistent with previous findings, in which recombinant SFRP2 was found to induce melanin synthesis in human primary melanocytes, via Wnt signaling activation, and to promote darkening of skin explants^18^.

Since *Sfrp2* promotes melanin synthesis in postnatal stages, we next asked whether expression levels of this gene differed between striped regions (i.e., stripes of different color). Using qPCR measurements from dissected P4 stripe skin regions, we found that *Sfrp2* was significantly lower in the light stripes (R2) compared to the flanking dark stripes (R1 and R3) (**Extended Data Fig. 6c**), an expression pattern drastically different from the dorsoventral gradient during mid-embryogenesis. Indeed, a time course analysis showed that *Sfrp2* expression in skin is temporally dynamic, shifting from a ‘high-low’ dorsoventral gradient at E16.5 to the ‘high-low-high’ pattern seen at P4 (**Extended Data Fig. 6c**). Taken together, our spatial expression data and *in vitro* functional experiments indicate that, during postnatal stages, *Sfrp2* undergoes a drastic shift in spatial expression and acts to promote melanogenesis via Wnt activation, thereby revealing an additional regulatory function during stripe pattern formation.

## Discussion

In birds and mammals, periodic pigment patterns arise from two sequential processes: the first occurs early in embryogenesis and specifies the positional boundaries of the future pattern (i.e.*, pattern establishment, or prepattern*); the second implements this prepattern through genes mediating pigment production during hair growth and cycling, giving rise to the observed color differences (i.e.*, pattern implementation*)^11, 43^. Here, we identify *Sfrp2* as a regulator of both the establishment of the pre-pattern and its implementation, a dual role that is made possible by its dynamic expression pattern throughout embryogenesis and early postnatal stages and its opposing effects on Wnt signaling (**Fig. 4g**). The involvement of *Sfrp2* at multiple stages of pattern formation implies a fine-tuned spatiotemporal regulatory control, likely achieved through a complex *cis*-regulatory architecture and the action of multiple binding factors^44^.

Secreted frizzled-related proteins (SFRPs) are known regulators of Wnt signaling that can exert their effects through a wide variety of mechanisms, including Wnt ligand sequestering, binding to Frizzled receptors, stabilization of Wnt-Frizzled complexes, and direct activation of Frizzled17. As such, SFRPs can either inhibit or activate Wnt signaling, and their effect is dependent on cell type, developmental stage, and cellular microenvironment^17^. In striped mice, embryonic expression of *Sfrp2* in a subset of dermal fibroblasts causes a decrease of epidermal CTNNB1 and LEF1, leading to alterations in placode number (**Fig. 4g**). Later in development, dermal papilla-specific expression of *Sfrp2* alters melanocyte behavior, causing an increase in the expression of Wnt targets and genes involved in melanogenesis (**Fig. 4g**). Thus, our results show that *Sfrp2* can lead to opposing regulation (i.e., activation and inhibition) of Wnt signaling within the same tissue.

It is noteworthy that elimination of *Sfrp2* led to a subtle, albeit consistent difference in the coat pattern of striped mice. This result is not surprising, as subtle changes in stripe width are consistent with predictions from our simulations. Moreover, it is likely that there are additional reinforcing/redundant mechanisms at play that act in concert to establish the dorsoventral orienting gradient and assure phenotypic robustness. For example, our bulk-level RNA-seq data from E16.5 embryos shows that there are additional Wnt inhibitors which are also expressed in a dorsoventral gradient, such as *Dkk2*, *Igfb4*, *Rspo2*, and *Rspo4* (**Extended Data Fig. 7**). Such molecules may act in coordination with *Sfrp2* to specify patterns of placode morphogenesis and are therefore still playing a substantial role in regulating this trait in *Sfrp2^-/-^* striped mice. Similarly, with respect to pigmentation, our previous work showed that hair color differences in striped mice are controlled by the interplay of melanocyte-autonomous (e.g., *Alx3*-mediated suppression of melanocyte differentiation) and non-autonomous processes (e.g., regulation of pigment-type switching by paracrine factors secreted from the dermal papilla, including *Asip* and *Edn3*)^10, 38^. Thus, although *Sfrp2* influences pigmentation, as our experiments show, it is clear that modulation of hair color in striped mice is achieved through the combinatorial effect of multiple genes^10^.

By integrating multidisciplinary approaches, our work reveals insights into the mechanisms by which spatial patterns are established in developing embryos. In addition, by extending *in vivo* genome editing approaches to wild-derived striped mice, a technique previously restricted to traditional mammalian laboratory models (i.e., laboratory mouse and rats), our work exemplifies the potential of using emerging model species to uncover the mechanistic basis of naturally evolved phenotypic traits and will open the door for functional studies in other mammals.

## Acknowledgments

We thank members of the Mallarino lab; Princeton LAR (Chris Dmytrow, Kirsten Gerhart, Grace Barnett, and Jamus McGuire) for help with striped mice husbandry; the LSI Genomics Core (Wei Wang, Jennifer M. Miller, Jessica Wiggins, and Jean Arley Volmar) for help with library preparation and sequencing; the Nikon Center of Excellence Confocal Microscopy Core (Gary Laevsky, Sha Wang); and members of the Rivera-Perez lab (Yeonsoo Yoon and Judith Gallant) for help with *in vivo* genome editing experiments. We also thank Eric F. Wieschaus, Girish Deshpande, and Philipp Holl for insights and discussion.

## Funding

This project was supported by an NIH grant to RM (R35GM133758). MRJ was supported by an NIH fellowship (F32 GM139253). SL was supported by a Presidential Postdoctoral Research fellowship (Princeton University). BB was supported by an NIH training grant (T32GM007388). CF was supported by an NIH fellowship (F32 GM139240-01). C.F.G-J. is partially supported by UC Irvine Chancellor’s ADVANCE Postdoctoral Fellowship Program. QN was partially supported by an NSF grant DMS1763272 and a Simons Foundation grant (594598).

## Author contributions

MRJ and RM conceived the project and designed experiments. MRJ performed all RNA sequencing experiments and bulk RNA sequencing analysis. SL performed the *in vitro* and *in vivo* genome editing in striped mice, with help from SAM and JAR-P. MRJ and SL performed all downstream processing and analysis of genome edited animals. PM and SYS did the mathematical modeling. CFG-J led the scRNAseq analysis, with support from MRJ and QN. MRJ and RM performed *in situ* hybridizations. MRJ, SAM, and BB performed the phenotypic characterization of striped mouse and laboratory mouse tissues, including immunofluorescence and histology. MRJ and SAM performed the melanocyte cell culture experiments. CF generated the “*Rhabdomyzed” Mus* genome and Liftover annotation. JG and AP generated the immortalized *Rhabdomys* fibroblasts. MRJ and RM wrote the manuscript with input from all authors.

## Competing interests

The authors of this publication declare that they have no competing interests.

## Data and materials availability

The bulk RNA-seq and scRNA-seq reads will be submitted under an NCBI BioProject.

## Methods

### Striped mouse husbandry

F10 descendants of wild-derived striped mice (*Rhabdomys pumilio)*, originating from Goegap Nature Reserve, South Africa, S 29° 41.56′, E 18° 1.60′, were originally obtained from a captive colony at the University of Zurich (Switzerland) and are maintained at Princeton University. *R. pumilio* are kept at a 16:8 light-dark cycle and given food *ad libitum*. All experiments performed were approved by Princeton University’s Institutional Animal Care and Use Committee.

### Embryo staging

To create a developmental timeseries of striped mouse embryonic development, embryos spanning mid-gestation were collected and ordered chronologically, based on embryo size and various morphological features including eye pigmentation, webbing of the forelimb and hindlimb, and skull morphology^45^. The stage at which epidermal hair placodes first become visible in striped mice was considered “stage matched” to *Mus* E15.5, the stage at which hair placodes first become visible by eye in *Mus*. Indeed, other morphological features in striped mice at this stage agreed remarkably well with that of E15.5 laboratory mouse embryos. This stage in striped mice was later confirmed to be E16.5 via a combination of vaginal cytology and ultrasound imaging. Staging of all striped mouse embryos was confirmed using vaginal cytology and ultrasound imaging.

### Sample preparation for in situ hybridization and immunofluorescence

Pregnant females of the appropriate stage were euthanized following approved protocols. Embryos were harvested, fixed in 4% paraformaldehyde (PFA), dehydrated in increasing concentrations of Methanol/PBS (25%, 50%, 75%, and 100%), and stored at −20°C. For whole-mount *in situ* hybridizations, embryos were rehydrated in decreasing concentrations of Methanol/PBT (75%, 50%, and 25%), and washed with PBS. For section *in situ* hybridizations and immunofluorescence, embryos were rehydrated as described above and then incubated in 10% sucrose overnight (O/N) at 4°C, followed by a 30% sucrose O/N incubation at 4°C. Embryos were then incubated in 50% Sucrose, 50% OCT (optimal cutting temperature) compound for 3 hours at room temperature, embedded in OCT, flash frozen, and cryosectioned (16um thickness) using a Leica CM3050S Cryostat. Slides were frozen at −80°C until use.

### Whole Mount in situ hybridizations

mRNA sequences for target genes were obtained either from NCBI (*Mus musculus*) or a denovo transcriptome (*R. pumilio*)^10^. For *R. pumilio*, we generated the following anti-sense riboprobes: *Dkk4* (543bp), *Wif1* (563bp), *Bmp4* (619bp), *Wnt10b* (360bp), *Dkk1* (546bp), and *Ctnnb1* (400bp). For *M. musculus*, we generated a *Dkk4* probe (550bp). All primer sequences are reported in Extended Data Table 2. All probes were sequence-confirmed using Sanger Sequencing. Whole-mount *in situ* hybridizations were then performed using previously described protocols^46^. Briefly, embryos were post-fixed with 4% PFA in PBS, washed with PBS, treated with 20ug/mL proteinase in PBT for 45 min, and incubated O/N with riboprobes at 65°C. The following morning, probes were washed with MABT (Maleic acid, NaCl, Tween-20) and incubated O/N with secondary anti-DIG antibody (1:2000) diluted in MABT, 2% Boehringer Blocking Reagent, and 20% heat-treated sheep serum. After washing several times with MABT, the signal was developed by incubating with NBT/BCIP. Once signal had developed sufficiently, the reaction was stopped by washing several times with PBT and fixing in 4% PFA O/N. Embryos were visualized using a SMZ18 stereo microscope (Nikon). At least 3 embryos per probe per species were analyzed.

### Bulk-level RNA sequencing

E16.5 striped mouse embryos (N = 3) were harvested from pregnant females and placed in 1x PBS on ice. Regions R1 (dorsalmost, placode barren), R2 (dorsalmost, placode rich), R3 (ventralmost, placode barren), and R4 (ventralmost, placode rich) were micro-dissected using morphological placodes as reference points. Regions from both sides of the midline were combined into a single tube containing 500ul of RNALater (Invitrogen) and stored at −20°C until RNA extraction. RNA was extracted using the RNeasy fibrous tissue mini kit (QIAGEN) per the manufacturer’s protocol. RNA-sequencing libraries were prepped using the TruSeq RNA Library Prep kit v2 (Illumina) and sequenced on a NovaSeq 6000 (2 x 65bp, paired-end). Pairwise differential expression analyses between the transcriptomes of dorsal skin regions R1-R4 were performed using DeSeq2 v1.30.1 from BioConductor (https://bioconductor.org/)^47^. Only genes differentially expressed with an adjusted p-value < 0.05 were considered. For visualization of all comparisons simultaneously, UpSet plots were generated using Upset R^48^.

### Modeling stripe patterns of placode formation

#### Model Preliminaries

All simulations in our numerical study were carried out on a domain of [−1, 1]. Throughout, we used values of *D_M_* = 1, *k_M_* = 1, *l* = 0.05, and *M_0_* = 1. Simulations were performed using the Dedalus package for spectral pde solution with a discretization of 512 modes^49^. Steady state solutions in each case were calculated via a variant of Newton’s method, and stability of stripes and uniform states were determined by calculation of the largest eigenvalues, using the methods previously described^50^ and a timestep of *dt* = 0.01.

### 3’-Single-cell RNA sequencing (scRNA-seq)

#### Sample preparation and library generation

Embryos from *R. pumilio and M. musculus* females were harvested at the relevant stages of pregnancy (i.e., E12.5-E15.5 for *M. musculus*; E13.5-E16.5 for *R. pumilio*). For all stages (except *R. pumilio* E16.5, see below), a single, rectangular piece of dorsal skin, symmetrical across the midline, was micro-dissected, using the forelimbs and hindlimbs as reference points to maintain consistency across stages and species. For *R. pumilio* E16.5 embryos, regions R1 (dorsalmost, placode barren), R2 (dorsalmost, placode rich), and R3 (ventralmost, placode barren) were micro-dissected using morphological placodes as markers and regions from both sides of the midline were combined. Dorsal skins from 3 embryos were pooled (with the exception of *M. musculus* E13, N=2) into a single tube containing 1mL 0.25% Trypsin/PBS. Skins were minced with fine scissors and transferred to an Eppendorf tube containing 10 mL of 0.25% Trypsin/PBS. Tubes were incubated at 37°C in a hybridization oven with gentle rotation for 15-30 minutes (timing was dependent on embryonic stage) until the tissue was well dissociated. Primary cell suspensions were passed through a 70um filter and diluted 1:5 with 4% Bovine calf serum (BCS)/PBS. Cells were centrifuged at 300 x g in a refrigerated centrifuge set at 4°C for 10 minutes. Supernatant was discarded, the cell pellet was gently washed with 4% BCS/PBS and resuspended in 100 µl of 0.1% Ultrapure Bovine Serum Albumin (BSA)/PBS (Invitrogen). Cell number and viability was assessed using trypan exclusion and a TC20 automated cell counter (BioRad). Single cell RNA-sequencing libraries were generated using the Chromium Single Cell 3’ Reagent Kits (v2) (10x Genomics) following the manufacturer’s instructions. Libraries were sequenced on an Illumina NovaSeq 6000 (25 x 125bp, paired-end)

#### R. pumilio annotation

To generate gene annotations for the striped mouse genome suitable for analysis of bulk RNA-Seq data, we transferred high-quality NCBI gene annotations from the laboratory mouse (Mus_musculus.GRCm38.100.chr.gff3) to the *de novo R. pumilio* assembly. To accomplish this, we used a homology-based lift-over procedure, implemented by the program LiftOff v1.4.1 using default parameters. Raw reads, genomes, and annotations have all been desposited under BioProject: PRJNA858857 BioSample: SAMN29758252.

#### Generation of a “Rhabdomyzed” genome

To both facilitate comparisons of transcriptomic data between *R. pumilio* and *M. musculus*, and to leverage the high quality gene annotations available for the latter, we generated an alternative, reference-guided assembly for the *R*. *pumilio* genome against which sequencing data was analyzed using previously described methods^51^. Briefly, genome reads for *R. pumilio* were aligned against the GRCm38 *M. musculus* genome using bwa-mem2 v2.1. Alignments were sorted and duplicates were removed with SortSam and MarkDuplicates from Picard tools (http://broadinstitute.github.io/picard/). Next, read pileups were generated using bcftools v1.11^52^ programs mpileup, ignoring indels to preserve the reference GRCm38 coordinate system. Base calls were then generated using bcftools call’s multiallelic caller. Finally, an initial reference-guided assembly was generated by projecting striped mouse base calls onto the reference mouse genome with bcftools consensus. This procedure was then repeated once more using the resulting consensus sequence from the first round as the new starting reference genome. Metrics using the *‘Rhabdomyzed’* genome compared favorably (e.g., number of cells identified, median genes identified per cell, and percentage of genes confidently mapped to transcriptome) to a *R*. *pumilio* assembly/Liftover annotation and was thus chosen for downstream analyses.

#### Processing of raw single-cell RNA-seq data

Raw FASTQ reads from *R. pumilio* samples were mapped with Cell Ranger (version 3.1.0) to the *Rhabdomyzed Mus genome* (see above) (Rhabdomys_to_Mus_bcftools_consensus3.fa/Mus_Musculus.GRCm38.101.gtf) Raw FASTQ reads from *M. musculus* samples were mapped with Cell Ranger (version 3.1.0) to the *M. musculus* mm10 reference genome (GRCm38.94.dna/GRCm38.84.gtf). Raw FASTQ reads will be deposited in BioProject.

#### Doublet/multiplet simulation and low-quality cell filtering

Raw count matrices from *R. pumilio* and *M. musculus* obtained from alignment with the *‘Rhabdomyzed’ Mus* genome *or* mm10 reference genome, respectively, were pre-processed and doublet/multiplets were simulated using Single-Cell Remover of Doublets (Scrublet)^53^ (version 0.2.1) with default parameters enabled. The doublet/multiplet score threshold was adjusted manually to intersect the bimodal distribution of the observed transcriptomes on the probability density histogram as suggested. Putative singlets were kept and used for downstream query and comparative analyses if and only if they met the following user-defined collective quality control metrics filtering criteria: a) cells with more than 350 but less than 5,000 genes were kept; b) cells containing less than 10% total mitochondrial gene content were kept; c) cells not identified as outliers falling outside a prediction confidence interval defined by a quadratic in a model of a gene vs. counts plot (P-value 1e^-3^) were kept^54^.

#### Anchoring, integration, and downstream analysis of single-cell RNA-seq data

We performed anchoring and integration of *R. pumilio* or *M. musculus* data sets using the Seurat package (Version 3.2.2, R Studio version 3.6.1)^55^. In brief, species-specific Seurat objects were created using individual, raw digitized count matrices as suggested by developer. Objects were merged and individual gene expression matrices were normalized, and variable genes/features identified. Data sets were anchored (n=30 dimensions) and integrated (n=30 dimensions). The integrated object was then scaled. Significant PCs used for clustering and neighbor-finding were identified using a combination of statistical and heuristic methods. Neighbors and clusters were identified with dimensions specified by user and visualized using two-dimensional t-distributed Stochastic Neighbor Embedding (tSNE) embedding. For all analysis, R1, R2, and R3 from *R. pumilio* E16.5 embryos were randomly down sampled to equal cell numbers before integration into a single object that was representative of E16.5 *R. pumilio* dorsal skin.

#### Cell type annotation

Cell types in *R. pumilio* and *M. musculus* were defined and annotated using a core set of *bona fide* E14.5 *M. musculus* gene biomarkers^22^ and are reported in Extended Data Table 1. Aggregate biomarker gene module scores were computed in Seurat. Aggregate biomarker gene module scores and individual gene expression profiles were Log-normalized and visualized as feature plots in two-dimensional tSNE embedding.

#### Quantification of Sfrp2^+^ fibroblasts

To quantify the number of *Sfrp2*^+^ fibroblasts in *R. pumilio* and *M. musculus*, we randomly sub-sampled *Sfrp2*^+^ fibroblasts (*Sfrp2*normalized counts > 0) in *R. pumilio* or *M. musculus* (n=25 iterations). To mitigate potential differences owed to sample size, we sub-sampled fibroblasts to the lowest number of total fibroblasts in anyone comparison. *Sfrp2*^+^ fibroblasts in *R. pumilio* or *M. musculus* were then normalized to total fibroblasts. Results from each iteration are plotted with confidence intervals and their correlation was determined with Spearman’s correlation. Results are represented as the average ratio of *Sfrp2*^+^ fibroblasts to total fibroblasts from 25 different iterations ± STDV.

#### Gene expression analysis

To quantify gene expression changes in *R. pumilio* fibroblasts that correlate with *Sfrp2* expression levels, we performed three sub-samplings of fibroblasts based on *Sfrp2* UMI counts: (1.) *Sfrp2-*high > 1.0; *Sfrp2*-low < 1, (2.) *Sfrp2-*high > 2; *Sfrp2*-low < 2, and (3.) *Sfrp2-*high > 3; *Sfrp2*-low < 3. Gene expression analysis was performed in Seurat (Wilcoxon Rank Sum test) to identify differentially expressed genes (DEGs, adjusted p-value < 0.05). Although good agreement was observed in all sub-samplings used, only genes identified as differential in all three sub-sampling were subjected to Gene Ontology analysis using Enrichr and significant GOs (Padj <0.05) were considered.

### In situ hybridization chain reaction (HCR)

A conserved region in the *Sfrp2* mRNA between *R. pumilio* and *M. musculus* was used to generate 16 probe binding sequences for *in situ* hybridization chain reaction (HCR). HCR was performed using the standard protocol for fixed frozen tissue sections available from Molecular Instruments®. High signal within limb sections of both species was used as a positive control.

### Immunofluorescence

Immuno-fluorescence was performed on tissue sections using standard procedures. Briefly, slides were washed with 1x PBS with 0.1% Tween (PBT) and blocked with 1x PBT/3% Bovine serum albumin (BSA) for 1 hour. Primary antibodies rabbit anti-LEF1 (Cell signaling #2230S, 1:100), rabbit anti-β-catenin (Sigma # C2206, 1:500), chicken anti-GFP (Novus Biologicals # NB100-1614, 1:200), or rabbit anti-Krt14 (BioLegend #905301, 1:1000) were diluted in 1x PBT/3%BSA and slides were incubated at 4°C O/N. The next morning, slides were washed several times with 1x PBT and secondary antibodies (goat anti-mouse Alexa Fluor 488, ThermoFisher, #A-21133) or (goat anti-mouse Alexa Fluor 546, ThermoFisher, #A-11001) were diluted 1:500 in 1x PBT/3%BSA and slides were incubated for 1 hour at room temperature. Slides were washed several times with 1x PBT, stained with DAPI to visualize nuclei, and mounted for imaging. Images were taken on a Nikon A1R confocal microscope. At least 3 embryos per condition were analyzed.

### Generation of Rosa^Sfrp^^2-GFP^ laboratory mice

*Srfp2* cDNA was synthesized (Azenta) and cloned into pR26 CAG/T2AGFP Asc (pR26 CAG/GFP Asc was a gift from Ralf Kuehn, Addgene plasmid # 74285, and modified to contain a T2A instead of IRES). The plasmid was inserted into the *Rosa26* locus by CRISPR-Cas9-mediated gene targeting. Cas9 (IDT) was complexed with an sgRNA (Millipore-Sigma) having the spacer sequence 5’-ACTCCAGTCTTTCTAGAAGA-3’. C57BL/6J zygotes were microinjected and transferred into pseudopregnant recipients. Founders were screened for the presence of GFP by PCR using primers GFP(F) and GFP(R)12 founders were further screened for 5’ end targeting using primers ROSA26J and SAR and for 3’ end targeting using GFP(F) and ROSA26L. All 12 were correctly targeted, 10 were chosen to confirm the presence of the cDNA by Sanger sequencing. Primer sequences are available in Extended Data Table 2.

### Quantification of hair follicle number

To examine the effect of dermal *Sfrp2* over-expression on hair follicle number, *Rosa^Sfrp^*^2^*^-GFP^*females were crossed to *Dermo1-Cre* (JAX stock #008712)^26, 27^ males and embryos were collected at E17.5. Embryos were fixed, embedded, and immunostained for GFP, as described above. GFP expression was mosaic, but a posterior dorsal region consistently showed high levels of GFP signal across all GFP (+) embryos. Within this region, follicle quantification was performed on equally sized areas of the skin after performing hematoxylin-eosin (H&E) stains to reliably identify hair follicles. In total, 4 double-transgenic and 4 control embryos were quantified. For each embryo, hair follicle number from 4 skin sections were counted and averaged with an equal sized region of the embryo. Statistical significance was assessed using an unpaired t-test. Statistical significance was assigned to p-values: p<0.05(*); p<0.01(**), p<0.001(***), p<0.0001(****).

### Quantitative Real Time PCR (qPCR)

Individual dorsal skin regions R1-R3 were dissected using either hair placodes (E16.5), hair length (E19.5) or pigmentation (P0, P4) as markers. Skin sections were stored in 500ul of RNALater (Invitrogen) and kept at −20°C until RNA extraction. RNA was extracted using the RNeasy fibrous tissue mini kit (QIAGEN) per the manufacturer’s protocol. cDNA synthesis was carried out using qScript cDNA SuperMix (Quantabio) and qPCR for *Sfrp2* was performed using SYBR Green qPCR Reagent (Quantabio) on a ViiA 7 Real-Time PCR machine (Applied Biosystems). Primers used for qPCR are listed in Extended Data Table 2.

### Over-expression of Sfrp2 in melanocytes

The LV-*GFP* plasmid used in this study has been described previously^56^. To generate LV-*SFRP2-GFP*, we amplified the *Sfrp2* coding sequence from an *Sfrp2* (NM_009144) Mouse Tagged ORF Clone (Origene CAT#: MR204070) using CloneAmp™ HiFi PCR Premix (Takara 639298) and cloned it (in frame) into the linearized LV-*GFP* plasmid using In-Fusion® Snap Assembly Master Mix (Takara #638947). High titer viruses were generated as previously described ^56^. Prior to transfection, MelanA cells (purchased from the Wellcome Trust Functional Genomics Cell Bank at St. George’s, University of London) were seeded in RPMI 1640 media supplemented with penicillin (100,000 U/L), streptomycin (100 mg/L), fetal bovine serum (10%), and tetradecanoyl phorbol acetate (TPA) (200nM). At ∼80% confluency, cells were centrifuged for 30min at 37°C and media was aspirated away and replenished with fresh media lacking antibiotics. LV-*GFP* or LV-*SFRP2*-*GFP* viral supernatant was added to each well along with a 1mg/mL polybrene diluted in bovine calf serum. Plates were gently swirled and incubated at 37°C/5% CO2 for 30 minutes. Plates were then centrifuged at 1100 x g for 30 minutes at 37°C. Media was removed, cells were rinsed twice with PBS, and fresh media was added. To enrich for cells with high transfection efficiency, cells with high GFP expression were sorted by fluorescence activated cell sorting (FACS) on a BD FACSAria II (BD Biosciences), grown to confluency, sorted one additional time, and grown to ∼80% confluency. Media was removed, cells were incubated in 500ul TriReagent (Zymo) for 5 minutes, flash frozen in liquid nitrogen, and stored at −80°C. RNA was isolated using a direct-zol RNA microprep kit (Zymo). cDNA synthesis was carried out using qScript cDNA SuperMix (Quantabio) and qPCR for *Mitf, Tyr, Axin2, C-myc*, and *Ccnd1* was performed using SYBR Green qPCR Reagent (Quantabio) on a ViiA 7 Real-Time PCR machine (Applied Biosystems), as described above. Primers used for qPCR are listed in Extended Data Table 2.

### In vivo genome editing in R. pumilio

#### Generation of immortalized R. pumilio fibroblasts

A skin biopsy was obtained from the flank of an adult male *R. pumilio* dermal fibroblasts were subsequently isolated following a previously published protocol^57^. In brief, hair, fat, and connective tissue were scraped away from the biopsy before digesting O/N at 4°C in HBSS without Ca^2+^ and Mg^2+^ (Thermo Fisher) containing dispase at a final concentration of 500 caseinolytic units/mL (Corning) and an antibiotic/antimycotic solution final concentrations of 100 μg/mL streptomycin, 100 IU/mL penicillin, and 250 ng/mL amphotericin B (HyClone). Following digestions, the epidermis was removed and discarded. The dermis was cut into small pieces, moistened with Dulbecco’s modified Eagle medium (Thermo Fisher; DMEM), and pressed into grooves scored into the wells of a 6-well tissue culture dish. The dermis was maintained in DMEM containing 10% (vol/vol) fetal bovine serum (Omega Scientific) and 1% (vol/vol) penicillin/streptomycin (Corning) at 37°C, 5% (vol/vol) CO2. Media was changed every 4–5 days and cultures monitored for fibroblast growth. Once sufficient outgrowth had occurred, the dermis was removed from the plate and the fibroblasts removed by trypsinization (0.05% trypsin-EDTA; Gibco) for expansion into larger culture dishes.

To generate the immortalized dermal fibroblast cell line, γ-retroviral pseudoparticles containing a transfer plasmid encoding simian virus 40 (SV40) large T antigen^58^ were produced in HEK293T cells. Cells were cultured on poly-L-lysine−coated 10 cm plates at 37°C, 5% (vol/vol) CO2 in 10% FBS DMEM. At ∼80% confluency, Xtremegene HP DNA transfection reagent (Roche Applied Science) was utilized per manufacturer’s directions to co-transfect the cells with 4 µg pBABE-neo-SV40 large T, a generous gift from Bob Weinberg (Addgene plasmid #1780; http://n2t.net/addgene:1780; RRID:Addgene_1780); 4 µg of a plasmid containing the genes for Moloney murine leukemia virus gag-pol; and 0.57 µg of a plasmid containing the gene for the G envelope protein of vesicular stomatitis virus. Supernatants were harvested 24, 48, and 72 hr post-transfection, stored at 4°C, and then pooled before passing through a 0.45 µm membrane filter (Millipore). Polybrene (Sigma-Aldrich; final concentration, 4 μg/mL) and HEPES (Gibco; final concentration, 2 mM) were added to the filtered supernatants; aliquots were prepared and at −80°C until needed. Primary dermal fibroblasts were seeded in 6-well plates for transduction so that cell confluency was 30–40% at the time of transduction. The cells were “spinoculated” in a centrifuge at 37°C, 931 rcf for 2 h with 2 mL of thawed, undiluted γ-retroviral pseudoparticles per well. The cells were subsequently kept at 37°C, 5% (vol/vol) CO2 and the media replaced with 10% FBS DMEM 6 hours post-spinoculation. The transduced cells were pooled once they achieved ∼80% confluency in the 6-well plate and subsequently expanded to prepare immortalized cell stocks. Cells were verified as negative for mycoplasma by testing with the MycoAlert Mycoplasma Detection Assay kit (Lonza) per the manufacturer’s instructions.

#### Guide-RNA design and testing

We designed 9 CRISPR sgRNAs targeting the *R. pumilio Sfrp2* exon1 using CRISPOR^59^. We then cloned each guide into the rAAV Nme2*Cas9* plasmid^36^ by replacing sg*Tyr* with sg*Sfrp2*^34^. Next, we transfected each AAV.Nme2*Cas9*.sg*Sfrp2* into immortalized striped mouse immortalized fibroblasts using lipofectamine3000 (L300008, Thermo Fisher). ∼3-4 days post transfection, we extracted genomic DNA using the Zymo Quick DNA Miniprep Plus Kit (Zymo) and performed targeted PCR on *Sfrp2* (primers are listed in Table S4). We then performed a T7E1 nuclease assay (M0302, New England Biolabs) according to the manufacturer’s protocol and resolved digested products on a 1.2% agarose gel containing STBY^TM^ safe DNA gel stain (Invitrogen) to assess excision efficiency of each designed sgRNA.

#### rAAV production, in vivo transduction, and genotyping

rAAV6. Nme2*Cas9*.sg*Sfrp2* was produced, concentrated, and purified by the PNI viral core facility at Princeton University. Female *R*. *pumilio* between 4 and 6 months of age were chosen to breed with age matched males. The presence of vaginal mating plugs or sperm was designated as days post coitum (d.p.c) 0.5. Pregnant females at d.p.c. 0.75 (N = 21) were anesthetized, administered with analgesics, and aseptically prepared for survival surgery. To deliver rAAV, we slowly injected 0.5-1 *μ*L rAAV (about 1∼3×10^9^ genome copy (GC)) mixed with Chicago Sky Blue dye (0.1%, Fisher Scientific, Cat # AAA1424214) into the oviduct ampulla using a glass micropipette with tip diameter of ∼10-30 *μ*m. The injection technique in *R*. *pumilio* was optimized based on previously described procedures in *M. musculus*^34, 60^. Operated animals were allowed to carry to term and pups were genotyped as follows: We collected ear punch tissue from individual animals and extracted genomic DNA using a Zymo Quick DNA Miniprep Plus Kit (Zymo). Next, we PCR amplified the genomic region containing the guide targeting site (primers are listed on Table S4) and resolved PCR amplicons using agarose gel electrophoresis. For F0 striped mice, PCR amplicons were further cloned into a pSC-amp/kan vector using StrataClone PCR cloning kit (240205, Agilent Technologies). We picked ∼20-50 single colonies per sample to determine the presence, composition, and frequency of targeted mutations via Sanger sequencing.

For knocking out *Tyr* in *R. pumilio*, 29 females were injected, 43 pups were obtained, and 9 of them carried gene edits in *Tyr*. Of these, we obtained one albino pup of homozygous knockout (*Tyr*^-/-^) (Fig.6B). 6 F0 founders were mated to wild-type animals to obtain germline transmission. Of these 6 founders, all of them successfully produced heterozygous animals.

For knocking out *Sfrp2* in *R. pumilio*, 21 females were injected, 57 pups were obtained, and 14 of them carried gene edits in *Sfrp2*. Of these, 11 F0 founders were mated to wild-type animals to obtain germline transmission. We obtained a total of 4 mutated alleles (2bp deletion, 13bp deletion, 466bp deletion, and 527 bp deletion); all of which produce a frameshift. 10 out of 11 founders tested successfully produced heterozygous F1 animals. F1 heterozygous striped mice containing the same mutated alleles were crossed to produce F2 animals. Genotypes of all F2 animals, determined by a combination of PCR, TOPO cloning, and Sanger sequencing, showed a mendelian ratio distribution (WT: N = 30; Heterozygous: N = 65; Homozygous: N = 27). At least two independently generated null alleles were tested in phenotypic analyses. Our designation of *Sfrp2^-/-^* mice encompasses animals that either share the same mutated allele or have a combination of two different mutated alleles.

#### Western blots

We collected fresh liver tissues of euthanized animals and prepared protein lysates in RIPA (50 mM Tris-HCl pH 8.0, 150 mM NaCl, 0.1%SDS, 0.5% Sodium deoxycholate, 0.1% Triton-X, 1 mM EDTA) supplemented with protease inhibitor cocktail (ab271306, Abcam). Protein concentration was quantified by Pierce Detergent Compatible Bradford Assay (Thermal Scientific). For SDS-PAGE, we loaded 20 *μg* of total protein per sample. Membranes were probed with primary antibodies at 4℃ O/N, washed with TBST 4x, and incubated in secondary antibody for 1 hour at room temperature. Primary antibodies: 1:100 *Sfrp2* (ab137560, Abcam, Cambridge, UK), 1:1000 *β*-tubulin (mAb #86298, Cell Signaling). Secondary antibodies: 1:20000 IRDye® 800CW Goat anti-Rabbit IgG (926-32211, LI-COR Biosciences), 1:20000 IRDye 680RD Goat anti-Mouse IgG system (926-68070, LI-COR Biosciences). Membranes were developed using an Odyssey CLx imaging system (LI-COR Biosciences).

### Quantification of Sfrp2^-/-^ phenotypes

#### Stripe width

Dorsal skin from equally staged P3 striped mouse *Sfrp2^-/-^* (N=7) or *Sfrp2^+/+^* (N=10) pups was dissected and flat-mount skins were prepped. Images of pinned skins were taken within a uniform lighting box with an EIOS2000D camera (Canon) at equal magnification. To quantify stripe width, FIJI (ImageJ) was used to draw line segments spanning each region of interest. For each image, 10 horizontal lines were measured along the Anterior posterior axis of the skin and an average width was calculated. For R1, R2, and R3, this was performed on each side of the midline. To measure width of the “striped area”, 10 lines were drawn connecting the most ventral boundary of R3 on both sides and averaged. “Width (% of striped area)” was calculated by dividing the average width of each region (i.e., R1, R2, R3, and midline) by the average width of the stripe area. Statistical significance was assessed using unpaired t-tests. Statistical significance was assigned to p-values: p<0.05(*); p<0.01(**), p<0.001(***), p<0.0001(****).

#### Pigmentation

To quantify pigment differences, a single image of 6 *R. pumilio* adults (3 *Sfrp2^+/+^*, 3 *Sfrp2*^-/-^) was taken within a uniform lighting box with an EIOS2000D camera (Canon). Color correction was performed on the image using a Pixel Perfect color checker and Adobe Lightroom software. To quantify pigment differences across animals, a consistently sized region of interest (ROI) was drawn within each stripe region and mean gray value within the ROI was measured using Adobe Photoshop. For R1-R3, mean gray values were measured in 3 subregions within each stripe region, on each side of the midline (6 ROIs total) and averaged. For the midline, mean gray values were measured in 5 subregions within the midline and averaged. Statistical significance was assessed using unpaired t-tests. Statistical significance was assigned to p-values: p<0.05(*); p<0.01(**), p<0.001(***), p<0.0001(****).

## Supplementary Information

**Extended Data Fig. 1:**
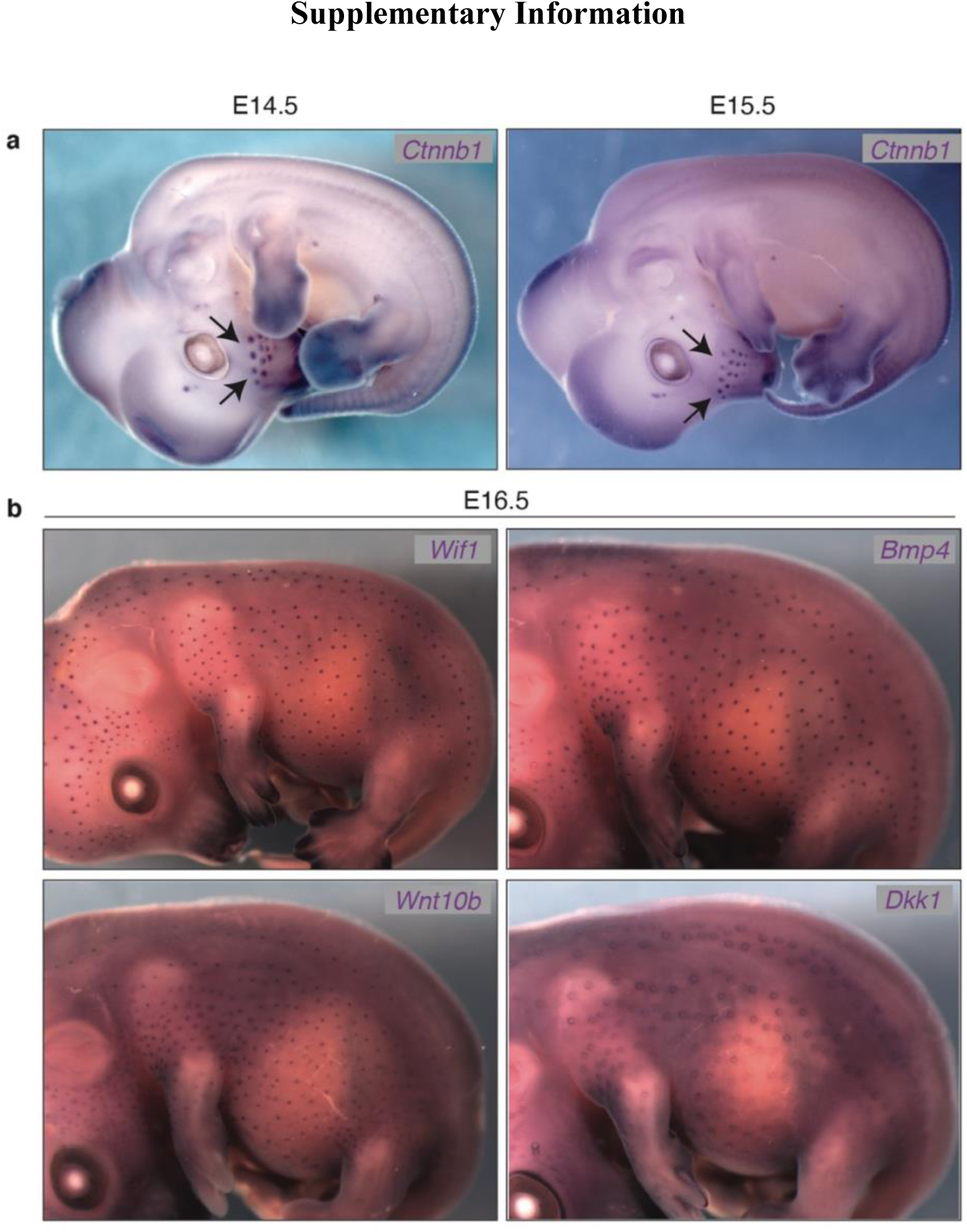
Patterns of hair placode formation in striped mice. **a**, Side views of E14.5 and E15.5 striped mouse embryos showing stages prior to the emergence of trunk hair placodes. Whole-mount *in situ* hybridization for an early placode marker, *Ctnnb1*, shows the presence of whisker placodes (arrows), which develop prior to trunk placodes. **b,** Side views of E16.5 striped mouse embryos displaying spatially restricted patterns of trunk hair placode formation, as visualized by whole-mount *in situ* hybridization for placode markers *Wif1*, *Bmp4*, *Wnt10b*, and *Dkk1*.

**Extended Data Fig. 2.**
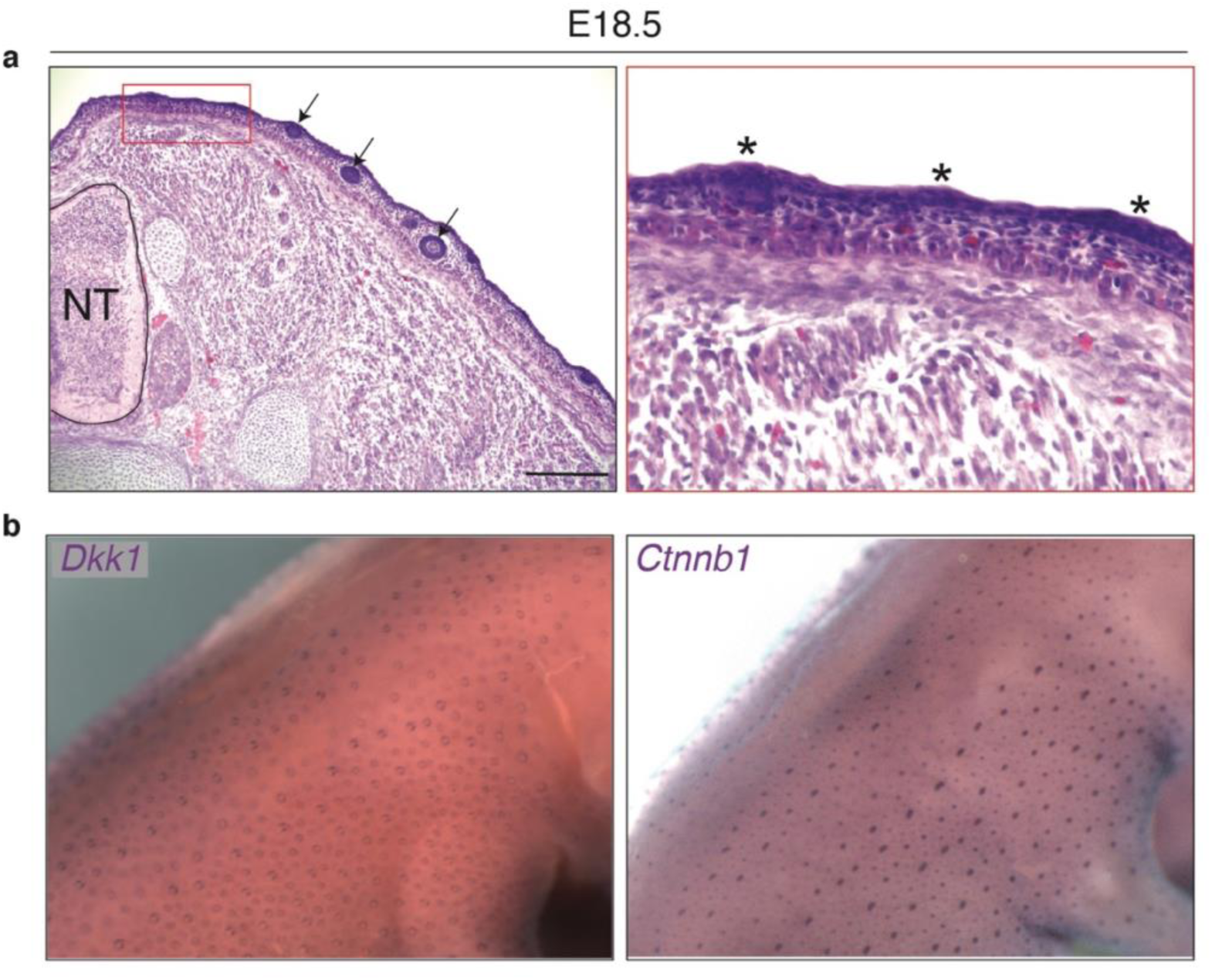
Placode formation during late embryogenesis. **a**, Hematoxylin-Eosin staining on cross sections of striped mouse E18.5 embryos reveals both mature placodes (arrows) and nascent placodes (asterisks); the latter are evidenced by thickening of the epidermis. **b**, Side views of E18.5 striped mouse embryos showing placode emergence in previously placode-barren regions, as visualized by whole-mount *in situ* hybridization for placode markers *Dkk1* and *Ctnnb1*. Scale bar in (**a**) = 100um.

**Extended Data Fig. 3.**
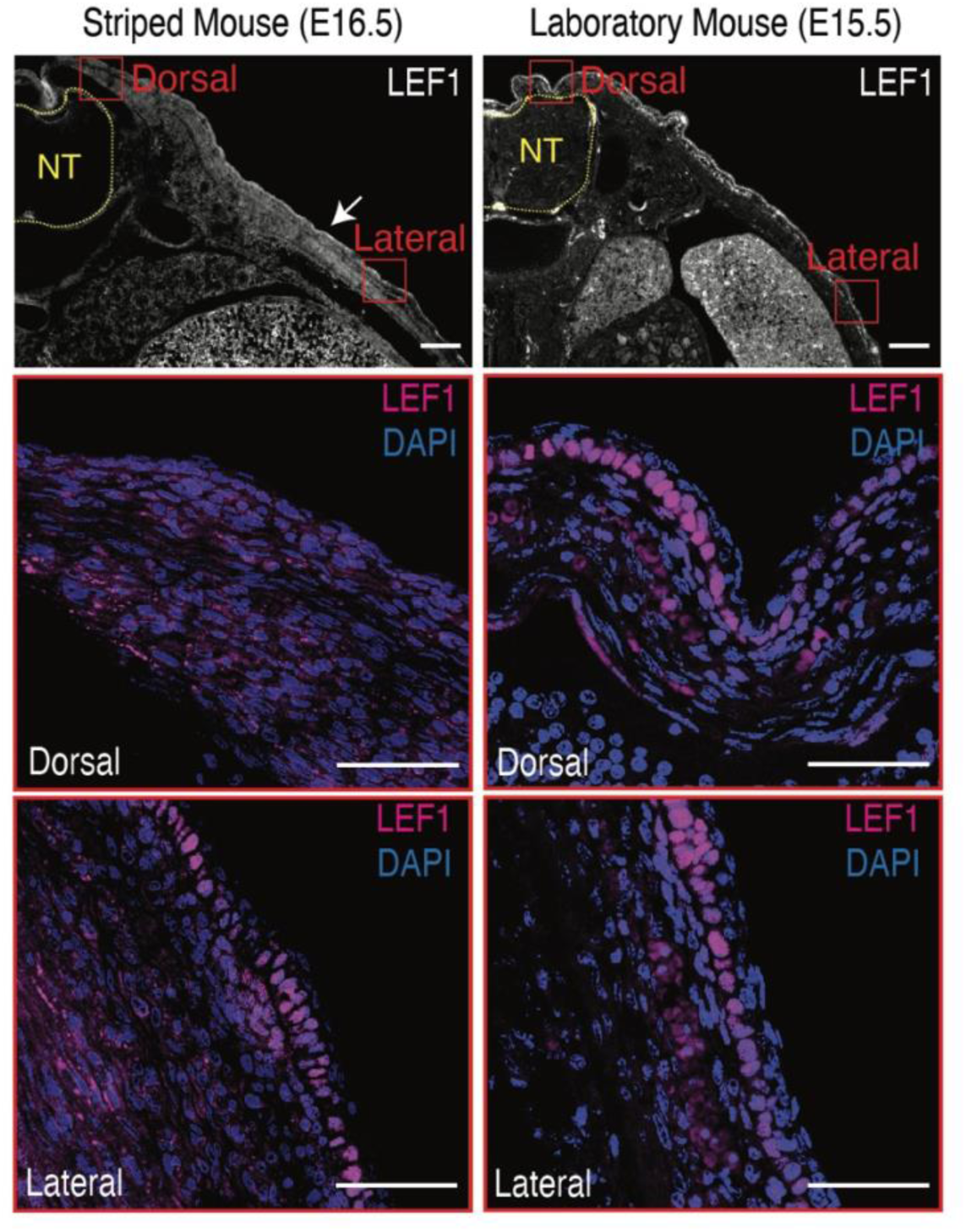
LEF1 immunostaining in staged matched striped (E16.5) and laboratory (E15.5) mouse embryos. IF for LEF1 in stage-matched striped mouse and laboratory mouse embryos. Red boxes denote zoomed-in regions. Scale bars: 200µm (zoomed out) and 50µm (zoomed in). NT = neural tube.

**Extended Data Fig. 4.**
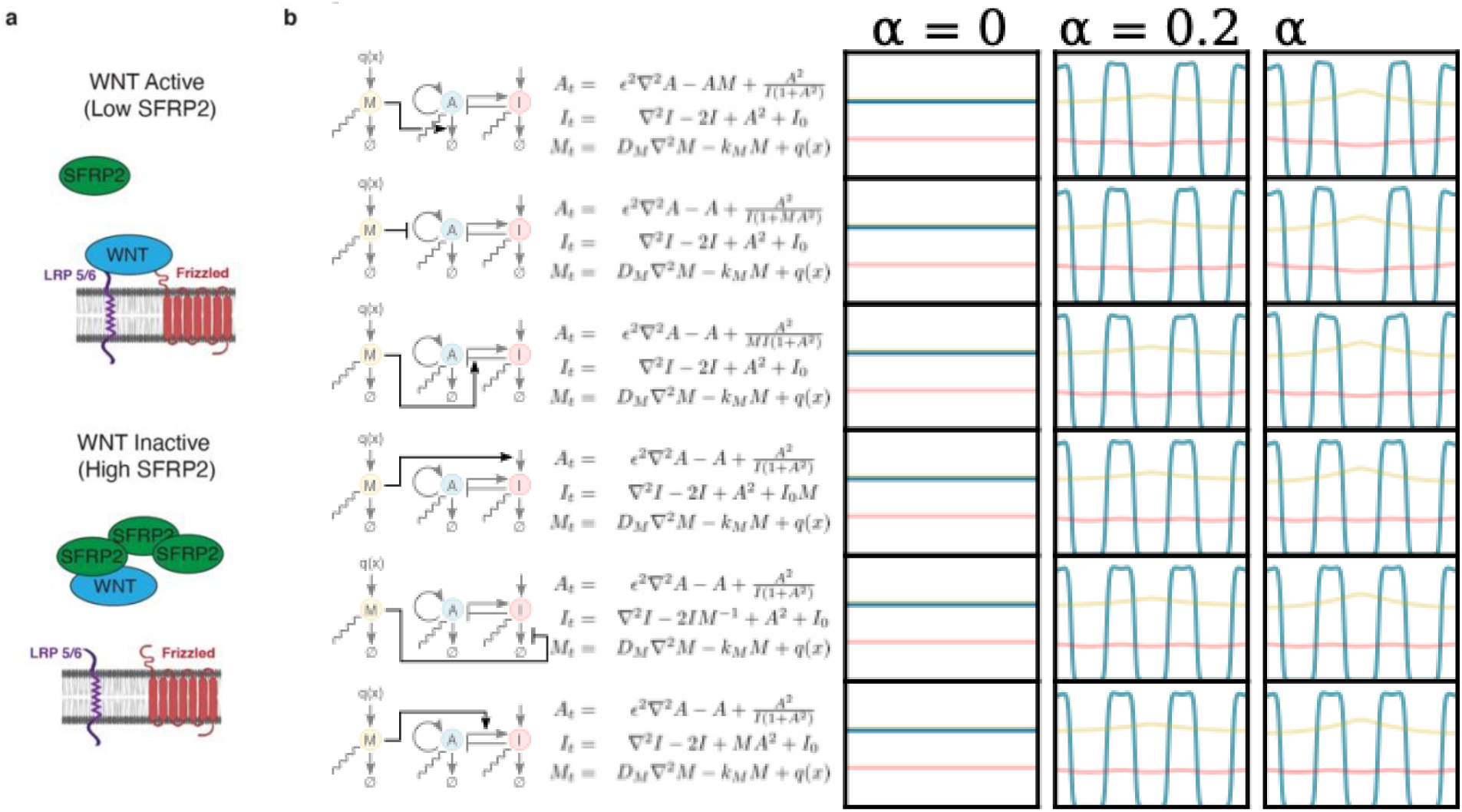
Gradient steepness increases central stripe width independent of model. **a**, Schematic showing the role of *Sfrp2* as an inhibitor of Wnt signaling. **b**, Each row depicts a schematic and equations governing a particular variant of our modulator-activator-inhibitor system (left) and the resulting simulations of stripe spacing for different gradient steepness values using these models (right). In all cases, gradient steepness affects stripe spacing.

**Extended Data Fig. 5.**
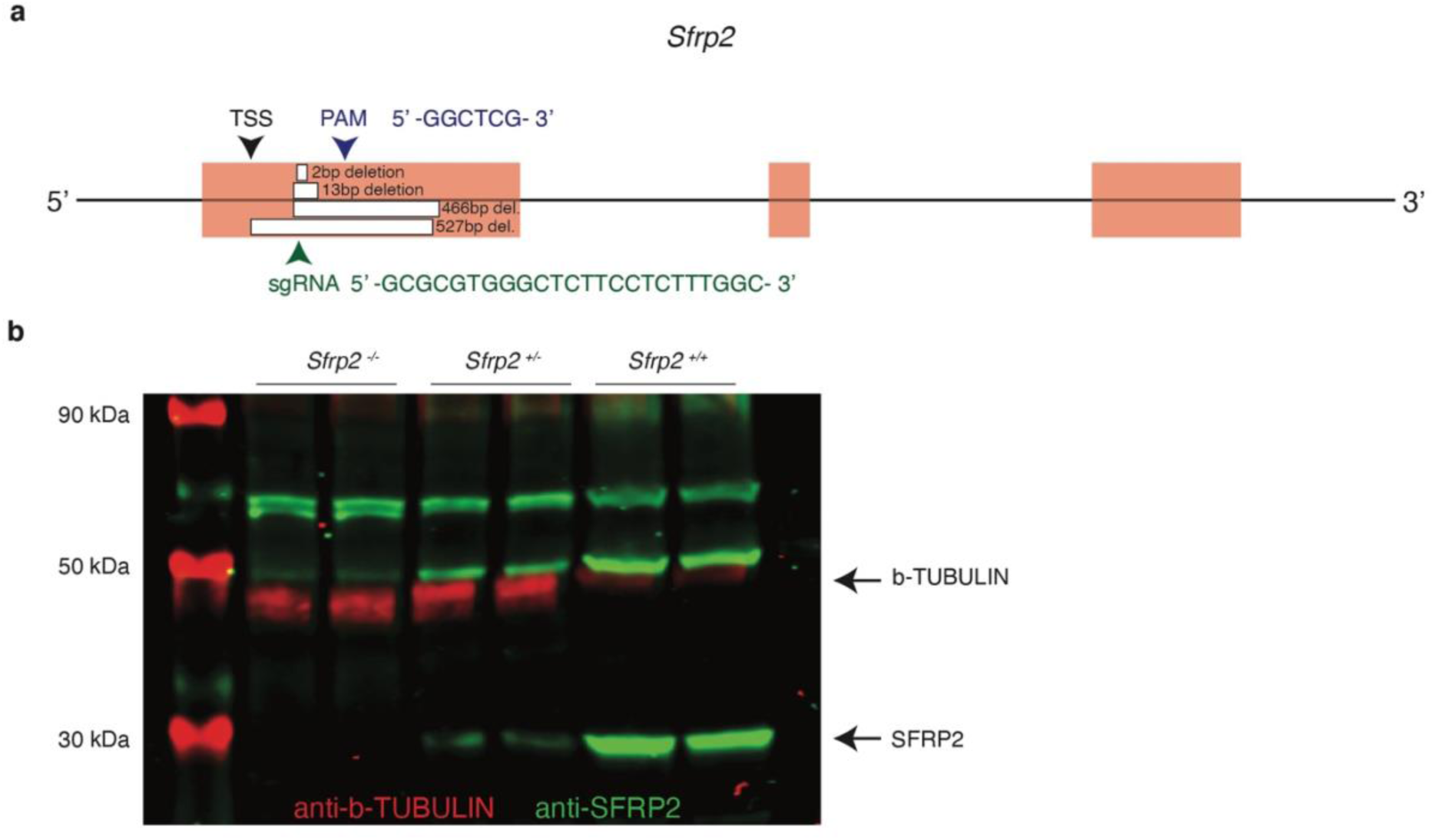
Generation of *in vivo* genome editing in striped mouse. **a**, Schematic of the *Sfrp2* locus (exons in red) showing the transcriptional start site (TSS), protospacer adjacent motif (PAM), and short guide-RNA (sgRNA) target/sequence. Four types of deletions were achieved: 2bp, 13bp, 466bp, and 527bp (white boxes). All mutations are predicted to cause frameshift mutations. **b**, Representative western blot of individuals carrying different combinations of wildtype and a 13bp deleted allele (wildtype: *Sfrp2^+/+^*; heterozygous: *Sfrp2^+/-^*; homozygous: (*Sfrp^-/-^*). *Sfrp^-/-^* have no detectable SFRP2 Protein (green). Bands ∼30 kDa correspond to SFRP2 protein. b-TUBULIN (∼50 kDa, red) was used as a loading control.

**Extended Data Fig. 6.**
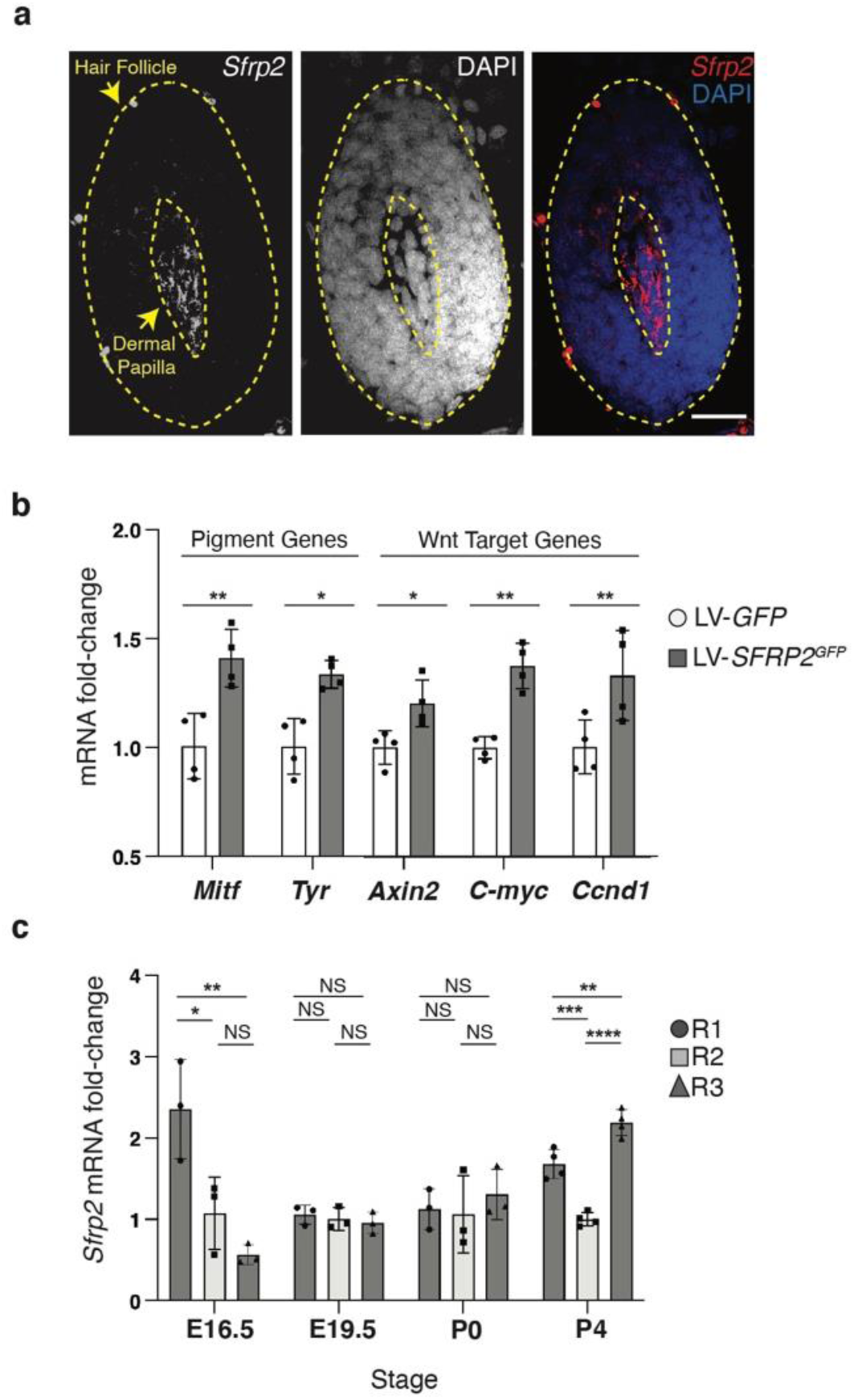
*Sfrp2* promotes melanogenesis by activating Wnt signaling. **a**, *In situ* hybridization showing specific *Sfrp2* expression in the dermal papilla of a P4 striped mouse hair follicle. **b**, Melanocytes were stably transduced with either a control (LV-GFP) or an experimental (LV-Sfrp2GFP) lentivirus and expression of Wnt targets and melanogenesis genes in stably transduced control and experimental cells, as determined was determined via qPCR. **c**, Quantitative PCR (qPCR) showing *Sfrp2* mRNA fold change levels along different dorsal skin regions in embryonic and postnatal stages. Statistical significance in panel (**b**) (** P < 0.01; *P < 0.05; N = 4) and in panel (**c**) (**** P < 0.0001; *** P < 0.001; ** P < 0.01; *P < 0.05, N = 3 for E16.5, E19.5, and P0, N = 4 for P4) was assessed using an ANOVA test.

**Extended Data Fig. 7.**
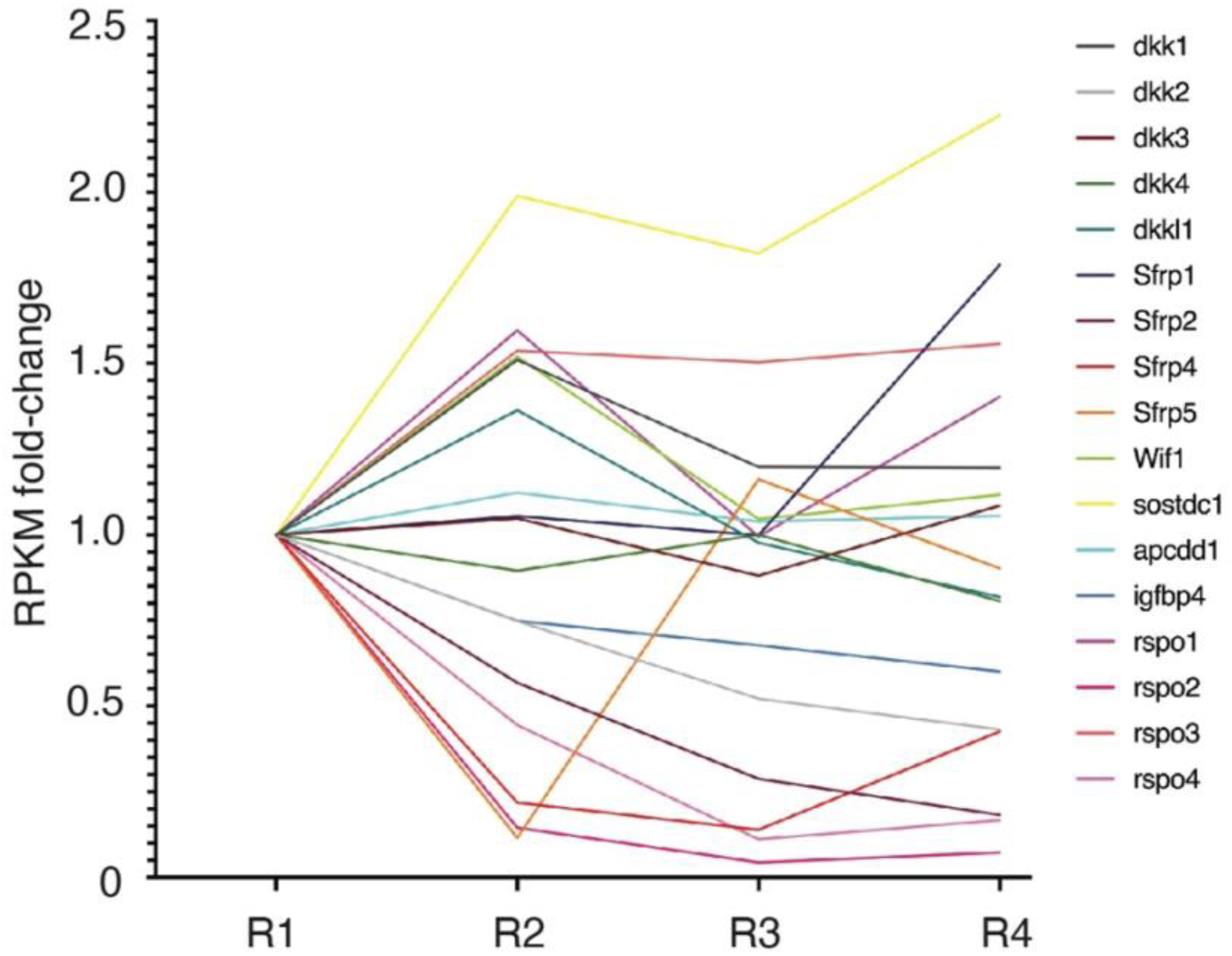
Expression of Wnt inhibitors in E16.5 striped mouse embryos. (A) Fold expression changes of Wnt inhibitor genes in skin regions (R1, R2, R3, R4) that were dissected for bulk RNA-seq analysis. Fold Expression changes were calculated from average FPKM values for each gene within each skin region. N=3.

**Extended Data Table 1.** Molecular markers used for cell type identification in scRNA-seq experiments.

| **Cell Population Identity** | **Markers used for identification** |
| --- | --- |
| Placode keratinocytes | *Lef1, Ptch1, Shh, Dkk4, Edar* |
| Basal Keratinocytes | *Krt14, Krt5* |
| Supra-basal keratinocyte | *Krt1, Krt10* |
| Granular keratinocytes | *Ivl, Lor* |
| Dermal Condensate | *Bmp7, Enpp2, Fgf10* |
| Fibroblasts | *Pdgfra, Col1a1, Lum, Dcn* |
| Satellite stem cells | *Pax7, Myf5* |
| Muscle | *Myog, Myod1* |
| Pericytes | *Rgs5, Pdgfrb* |
| Vascular endothelial | *Tie1, Pecam1* |
| Lymphatic | *Mpz, L1cam* |
| Immune | *Ptprc* |
| Melanocytes | *Tyr, Dct, Mlana* |
| Schwann | *Errb3, Mpz and L1cam.* |

**Extended Data Table 2:**
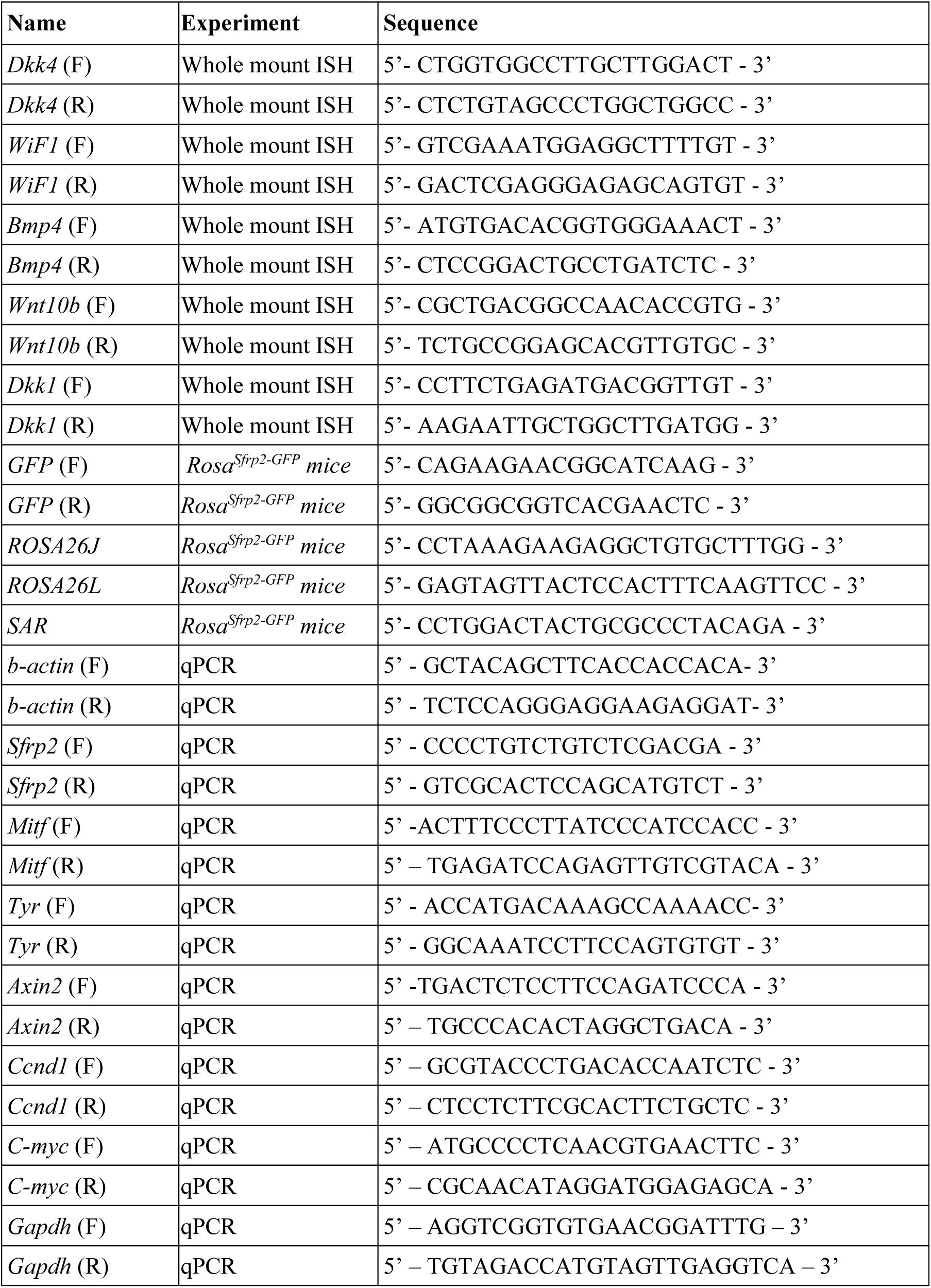
Primer sequences used in this study.

Supplementary Table 1: Differentially expressed genes between Region 1-4 of E16.5 striped mice skin. (Data_S1.xlsm)

Supplementary Table 2: Differentially expressed genes between Sfrp2^high^ and Sfrp2^low^ fibroblast populations. (Data_S2.xlsm)

## Notes

### Competing Interest Statement

The authors have declared no competing interest.

### Summary of Updates

The order in which the data are presented has been modified. The mathematical simulation is all part of the same figure

